# Insect hosts are nutritional landscapes navigated by fungal pathogens

**DOI:** 10.1101/2024.01.04.574030

**Authors:** Henrik H. De Fine Licht, Zsuzsanna Csontos, Piet Jan Domela Nijegaard Nielsen, Enzo Buhl Langkilde, August K. Kjærgård Hansen, Jonathan Z. Shik

**Author notes:** Corresponding author, Section for Organismal Biology; Department of Plant and Environmental Science; University of Copenhagen; 2000 Frederiksberg; Denmark.

## Abstract

Nutrition can mediate host-pathogen interactions indirectly when specific deficiencies (e.g. iron or glutamine) constrain host immune performance. Nutrition can also directly govern these interactions since invading pathogens colonize finite landscapes of nutritionally variable host tissues that must be optimally foraged during pathogen development. We first used a conceptual framework of nutritional niches to show that insect-pathogenic *Metarhizium* fungi navigate host landscapes where different tissues vary widely in (protein (P) and carbohydrates (C)). We next tested whether host-specific *Metarhizium* species have narrower fundamental nutritional niches (FNN) than host-generalists by measuring pathogen performance across an *in vitro* nutritional landscape simulating a within-host foraging environment. We then tested how developing pathogens navigate nutritional landscapes by developing a liquid-media approach to track pathogen intake of P and C over time. Host-specificity did not govern FNN dimensions as three tested *Metarhizium* species: 1) grew maximally across C treatments assuming P was present above a lower threshold, and 2) similarly initiated dispersal behaviors and sporulated when either C or P became depleted. However, specialist and generalist pathogens navigated nutritional landscapes differently. The host specialist (*M. acridum*) first prioritized C intake, but generalists (*M. anisopliae*, *M. robertsii*) prioritized P and C according to their availability. Numbers of known hosts may be insufficient to delimit pathogens as specialists or generalists since diverse hosts do not necessarily comprise diverse nutritional landscapes. Instead, immune responses of hosts and nutritional niche breadth of pathogens are likely co-equal evolutionary drivers of host specificity.

## INTRODUCTION

Pathogens obtain all nutrients from the infected host and nutrition can directly govern the outcome of host-pathogen interactions [1]. For an invading pathogen the host body represents an ecological habitat with nutrient compositions varying between host tissues and organs [2–7]. The breadth of nutrient compositions that support pathogen growth is termed the fundamental nutritional niche (FNN), and a developing pathogen can in principle deplete different body tissues to meet FNN needs specific to each pathogen growth stage [8]. During host colonization from pathogen entry, establishment, growth, and development, different types of nutrients can therefore be limiting. Single nutrients can be important for the outcome of host-pathogen interactions with the effect on the pathogen ultimately determined by the combined effects of nutrition and immune defenses [7,9]. Host-pathogen interactions can also be indirectly mediated by nutrition when specific deficiencies (e.g. iron [10,11] or glutamine [12,13]) constrain host immune performance [14–16], or the metabolic state of the host for example due to stressful conditions alter availability of nutrients for invading pathogens.

Insects are the most diverse lineage of multicellular organisms [17], which have contributed to the high diversity of insect pathogens such as entomopathogenic fungi [18]. For example the globally distributed diverse fungal genus *Metarhizium* contains more than 50 species [19]. These fungi span a continuum of host specificity from specialists (e.g. the locust-specific *M. acridum*) to generalists that infect most orders of insects (e.g. *M. anisopliae* and *M. robertsii*). This host-specificity continuum helps explain the diversity of pathogen life history strategies, with specialists tending to prioritize rapid growth during infection and generalists tending to initially be slower growing with higher investments in toxin production (e.g. destruxins) [20–22]. We hypothesized that this life history continuum mediates physiological allocation and nutritional metabolism such that host specificity also governs FNN dimensions.

Fungi regulate their metabolism according to available environmental nutrients via elaborate transcription regulatory programs [23]. For fungal pathogens, sensing and acquisition of different carbon and nitrogen sources mediate virulence by influencing secretion of fungal enzymes, cell wall remodeling, and morphological development [24,25]. After penetrating an insect host’s cuticle, an invasive *Metarhizium* fungus fuels population growth as single-celled yeast-like elements that initially consume the mostly C-rich hemolymph [26] (Fig. 1A). Over the ensuing days and coinciding with death of the host, foraging can shift to P-rich muscle and reproductive tissues [27] (Fig. 1A). This dietary switch enables the pathogen to meet its changing nutritional needs [28] associated with expanding its network of foraging hyphae. Ultimately, the fungus switches to production of asexual spores (conidia) that disperse from the nutritionally depleted host (Fig. 1A).

**Figure 1.**
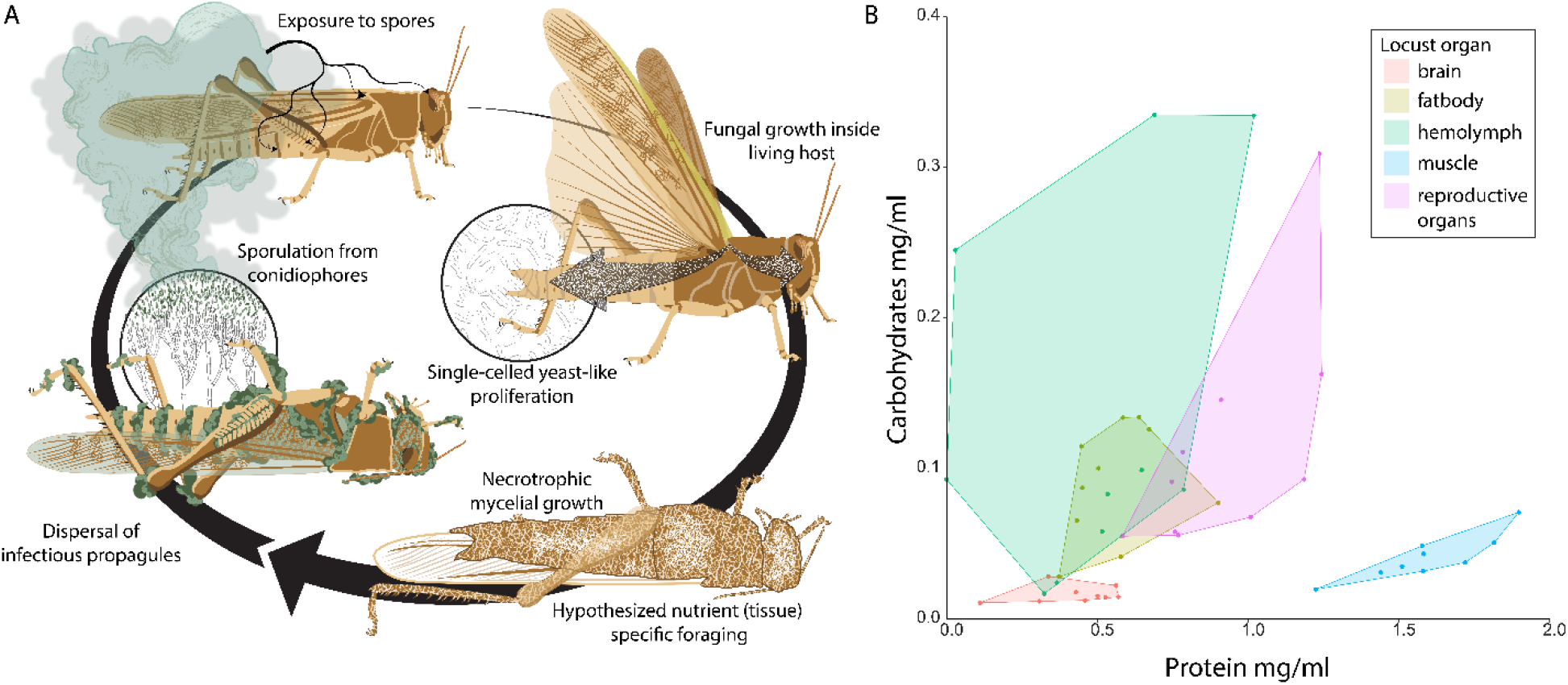
Visualizing a locust host as a landscape of nutritionally variable tissues foraged by developing *Metarhizium* fungal pathogens. **A.** Infection starts after the locust is exposed to *Metarhizium* conidia that adhere to its cuticle. The conidia then germinate to form appressorial cells that penetrate the cuticle using mechanical forces and enzymatic degradation. Inside the body cavity, single-celled ovoid hyphal-bodies called blastospores obtain nutrients and proliferate throughout the hemolymph. After one to several days, the insect dies and *Metarhizium* switches to nectrotrophic growth, which coincides with penetration of all body tissues. After accessible nutrients are depleted, the fungal pathogen initiates dispersal by producing green conidia that are visible emerging from the insect. **B.** We confirm that the different tissues comprising the foraging landscape withing the insect host contain different ratios and concentrations of protein and carbohydrates and thus provide opportunities for nutrient specific foraging and possibly niche partitioning among competing pathogen species. Each dot represents a different tissue sample.

We first tested the supposition that an insect body provides a finite landscape of nutritionally variable tissues by dissecting and measuring protein (P) and carbohydrate (C) content of individual organs (e.g., hemolymph, muscle, brain, fat body, reproductive organs). Secondly, we show that insect-pathogenic *Metarhizium* fungi can navigate host landscapes where different tissues vary widely in P and C. We specifically tested whether a host-specific *Metarhizium* species has a narrower fundamental nutritional niche (FNN) [23,24] than host-generalists by measuring pathogen performance across an *in vitro* nutritional landscape simulating a within-host foraging environment. Third. we tested how developing pathogens navigate nutritional landscapes by developing a liquid-media approach to track pathogen intake of P and C over time. Contrary to predictions, host-specificity did not govern FNN dimensions as three tested *Metarhizium* species: 1) grew maximally across C treatments assuming P was present above a lower threshold, and 2) similarly initiated dispersal behaviors and sporulated when either C or P became depleted. However, specialist and generalist pathogens navigated nutritional landscapes differently. The host specialist (*M. acridum*) first prioritized C intake, but generalists (*M. anisopliae*, *M. robertsii*) prioritized P and C according to their availability.

## RESULTS

### Mapping the nutritional landscape of tissue resources within a host insect

We first tested whether an insect body provides a finite landscape of nutritionally variable tissues (e.g., hemolymph, muscle, brain, fat body, reproductive organs). As predicted, these tissues have distinct blends of protein (P) (Kruskal-Wallis χ^2^= 32.801, df = 4, p < 0.0001) and carbohydrates (C) (Kruskal-Wallis χ^2^=27.353, df = 4, p < 0.0001) (Fig. 1B, Fig. S1). Muscle tissue has significantly higher P levels and brain tissue had lower C levels than other tissues (Fig. 1B, Fig S1). Hemolymph C levels varied more than other tissues but tended to be significantly higher than other tissues (Fig. 1B, Fig S1).

### Linking host-specificity and fundamental nutritional niche breadth

We next hypothesized that host specificity mediates a pathogen’s physiological needs since specialists tend to prioritize rapid growth during infection and generalists tend to prioritize toxin production (e.g., destruxins) as they grow[20,25,29]. These ideas can be extended beyond needs for individual nutrients to the FNN in multiple nutritional dimensions that account for life history variation. It is useful to conceptualize that generalist pathogens are ecologically similar to the most widespread invasive species whose broad FNNs can enable introduced propagules to establish in diverse nutritional landscapes across many ecosystems[1,30,31].

Fungal pathogens in the genus *Metarhizium* are ideally suited to test hypotheses linking host-specificity, FNN breadth, and ontogenetic shifts in nutrient intake. Here, we predicted the host specialist species *M.* acridum that only infects locusts will have narrower FNN dimensions than two host generalist species *M. anisopliae* and *M. robertsii* that infect most orders of insects (Fig. 2A). We used an *in vitro* approach[32] to quantify the nutritional blends that maximize pathogen growth performance (radial area) and fitness (timing and amount of spore production) (Fig 2B). By confining fungal isolates to 36 nutritionally-defined media treatments that systematically varied ratios and concentrations of P and C, we simulated nutritional landscapes within insect hosts where different tissues provide an array of nutritional foraging options (Fig. 2C).

**Figure 2.**
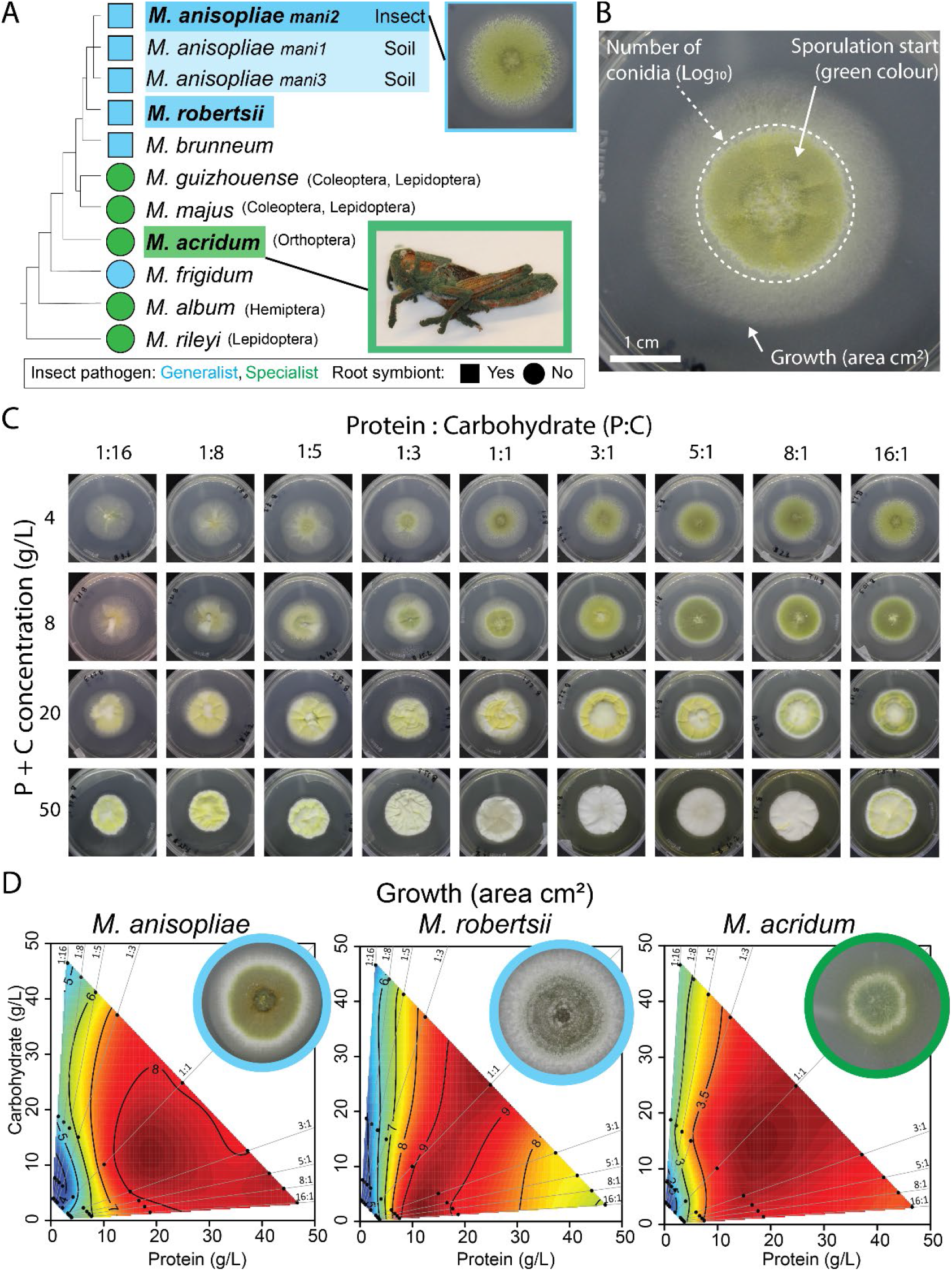
Linking host specificity with fundamental nutritional niche (FNN) breadth across species of *Metarhizium* pathogens. **A.** We used a schematic phylogeny of the fungal genus *Metarhizium* to map host-specificity and potential for free-living existence in the plant rhizosphere. Species analysed in the present study are marked in bold font. The species *M. anisopliae* is currently divided between three distinct phylogenetic groups where *mani3* mostly infects insects while *mani1* and *mani2* are mostly isolated from soil. **B.** We measured three performance traits to assess FNN dimensions. We illustrate these traits with an *in vitro* image of *M. anisopliae* (ESALQ1604) cultivated on a 1:1 Protein:Carbohydrate agar media at 4 g/L P + C. Mycelial growth was measured as the area (mm^2^) of the mycelium. Reproductive effort was measured in terms of total number of green conidia and the onset day of green coloration indicating the switch to reproduction. **C.** We quantified variation in pathogen performance across an *in vitro* nutritional landscape using fundamental nutritional niches (FNNs). FNN heatmaps were based on measures of pathogen performance recorded across 36 nutritionally-defined media treatments varying systematically in P:C ratios and P + C concentrations. **D.** FNN heatmaps of fungal growth area (cm^2^) are visualized for *M. anisopliae, M. robertsii,* and *M. acridum* using mean values of three isolates. Red colors indicate high growth values and blue colour depicts low growth values. Statistical analyses used to support the interpretations of heatmaps are provided in Table S1 (isolate-level analyses) and Table S2 (species level analyses).

In contrast to the host specificity hypothesis, all *Metarhizium* species studied had similarly broad FNNs for hyphal growth area, performing maximally across most provided blends of P and C (Fig. 2D, Table S1, Table S2). Yet, growth performance also exhibited a nutrient-specific response to the most imbalanced diets. P limitation (< 5 g/L) strongly constrained hyphal growth for each species even when C was available (p < 0.0001, Table S2), but C limitation did not constrain hyphal growth when P was available (Fig. 2D, Table S2). Thus, while early-infection success hinges on consumption of C-rich hemolymph by single-cell yeast-like propagules, the transition to threadlike hyphae at intermediate growth stages likely depends on the pathogen’s perfusion of increasingly P-rich tissues (e.g., muscle). Accelerated decomposition of tissues following host death likely facilitates this tissue perfusion and thus the fungal shift to necrotrophic growth.

Fungi have elaborate transcription regulatory programs for capitalizing on available nutrients[33]. The metabolic pathways underlying fungal pathogen virulence likely codify these links between specific nutrients and the secretion of enzymes and small effector proteins, the remodeling of cell walls, and shifts in morphological development[28,34]. In this way, tissue-specific foraging timelines of gradually depleted host landscapes are likely optimized by natural selection to target nutrients that trigger physiological shift from hyphal growth to spore production (Fig. 1A). To test this hypothesis, we next used FNNs to link the timing and magnitude of spore production with depletion of specific P:C ratios and P+C concentrations.

Nutritional co-limitation (low concentrations of both P and C) tended to trigger the start of sporulation six days earlier than when P and C were both abundant (i.e., red areas in the lower left corners of heatmaps spanning all P:C ratios (Fig. 3A, Table S1). In contrast, total spore number was maximized when only C was limiting (horizontal red area) or only P was limiting (vertical red area) resulting in ‘L-shaped’ FNNs (Fig. 3B, Table S1). These results move beyond observations that starvation conditions in depleted cadavers can increase the virulence[21,35] and number[36] of *Metarhizium* spores to show that multiple nutrients in isolation or in combination can mediate variation in pathogen reproductive effort.

**Figure 3.**
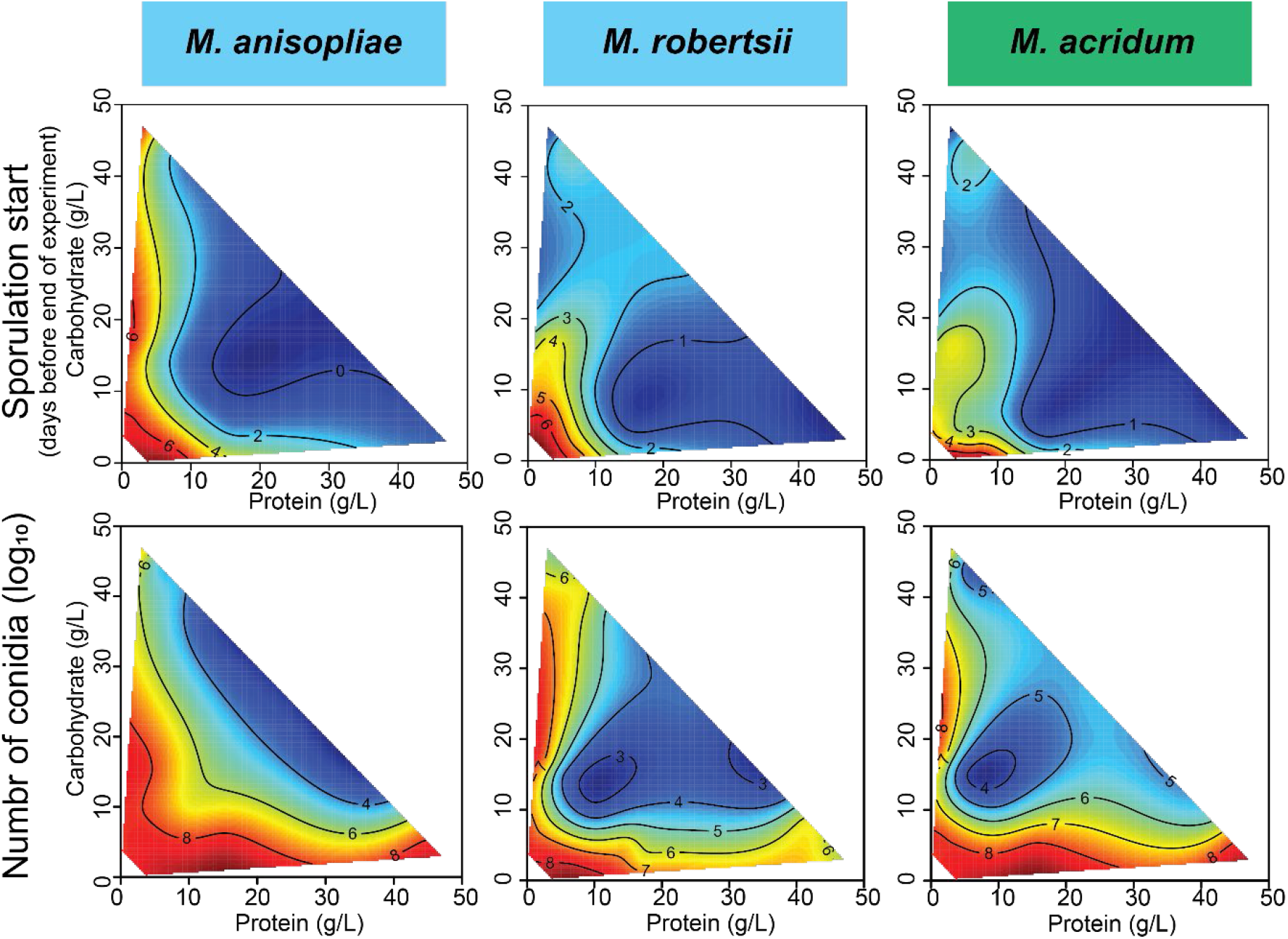
Comparing FNNs for reproductive effort traits across three species of *Metarhizium.* Heatmaps show how onset day of sporulation (top panel) and Log_10_(number of conidia) (bottom panel) vary across the 36 nutritionally-defined media treatments shown in Fig. 2C during the 11 days experiment. The heatmaps show the average values of three isolates of each of the three species and red heatmap colour show early sporulation (top panel) or high numbers of conidia (bottom panel) and blue heatmap color depicts low values of these traits. Statistical analyses used to support the interpretations of heatmaps are provided in Table S1 (isolate-level analyses) and Table S2 (species level analyses).

This nutritional niche framework can also help parse community-level dynamics among invading pathogen species and other microbial symbionts that face tradeoffs in their allocation to growth or defensive toxins. First, the need to inhibit other microbes may be reduced if pathogen species ecologically partition host tissues based on their nutritional composition. Second, the physiological costs of persisting on available but nutritionally suboptimal tissues may be low since each of the *Metarhizium* pathogens we studied could maximize spore numbers even when nutrient limitation induced earlier sporulation (Fig. 3).

### Do fungal pathogens selectively prioritize nutrients when navigating host landscapes?

We next analyzed the foraging behaviors of developing fungal pathogens by creating a glass-bead[37] liquid-media[38] approach to measure their P and C intake over time. We further simulated different nutritional landscape ‘starting points’ representing different potential host species by confining fungi to Petri dishes with different P:C ratios (1:3 or 3:1) and different P + C concentrations (15 g/L or 50 g/L). By repeatedly sampling small amounts of liquid media from each Petri dish over six days, we tested whether fungi preferentially depleted P or C and whether this depletion compensated for an initial P:C ratio imbalance or P + C concentration deficit (Fig. 4A).

**Figure 4.**
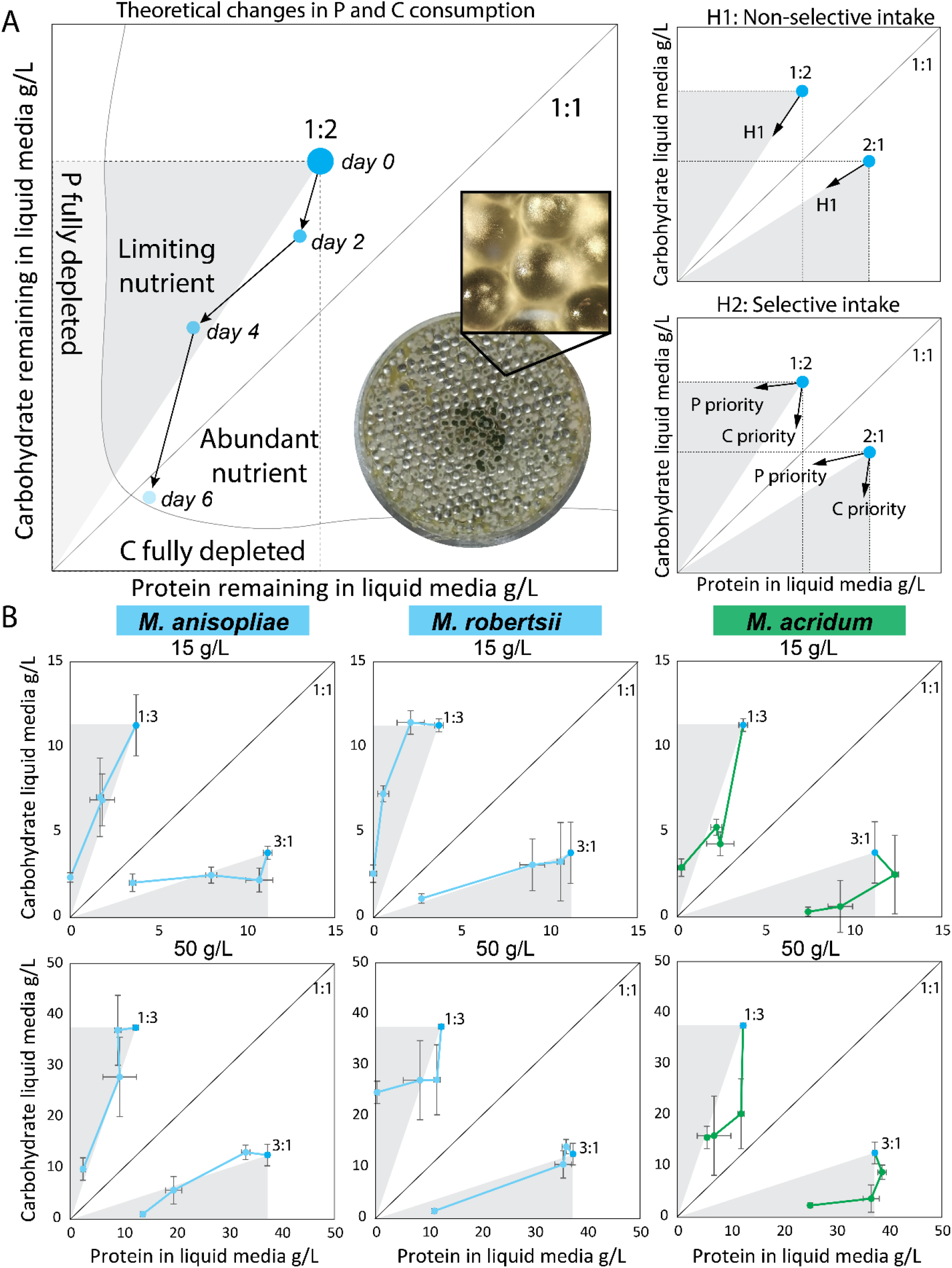
Comparing nutritional foraging strategies of generalist and specialist *Metarhizium* pathogens. **A.** Nutrient foraging strategies were measured *in vitro* by measuring the amount of protein and carbohydrate remaining in nutritionally-defined liquid media after two, four, and six days of fungal growth. As an example, we show fungal foraging from a 1:2 P:C starting point. The dashed line rectangle shows the nutritional geometric area available to the fungus for nutrient consumption. The grey triangle shows the limiting nutrient (P) and the white triangle shows the abundant nutrient (C). The non-selective intake hypothesis is illustrated by these diagonal segmented arrows showing that daily fungal P and C intake leads to the origin in the lower left corner. Here, P and C are both totally and passively depleted based only their relative abundance in the 1:2 P:C growth media. In contrast, the selective intake hypothesis shows either horizontal arrows leading to the Y axis (selective P foraging) or vertical arrows leading to the X axis (selective C foraging). Here, intake is independent of the initial relative abundance of either nutrient in the growth media. Picture shows *M. acridum* growing in petri-dish filled with glass beads and liquid media with a close-up showing how the glass beads support fungal growth. **B.** Nutrient foraging in three species of *Metarhizium* measured in a factorial combination of P:C ratios (1:3, 3:1) and P + C concentrations (15 g/L in the top row, 50 g/L in the bottom row). Each individual dot is the mean ± SE of three measured replicates.

If fungi do not selectively prioritize P or C, we predicted that their intake would passively track the provided P:C ratio, and their daily nutrient intake arrows would follow a diagonal line from the starting point to origin (Fig. 4A). If fungi selectively prioritize P or C, daily intake arrows would form a horizontal line from the starting point to the Y axis (P preferentially depleted) or a vertical line from the starting point to the X axis (C preferentially depleted) (Fig. 4A). Host-specificity appeared to govern nutrient-specific foraging strategies. Generalists *M. anisopliae* and *M. robertsii* tended to consume P and C in the same P:C ratios provided by liquid media (i.e. diagonal arrows, Table S3), while occasionally prioritizing P or C on some sampling days (Fig. 4B, Table S3). By non-selectively consuming nutrients at provisioned ratios, the generalists appear to exhibit flexible nutrient intake consistent with ability to invade diverse nutritional landscapes found within diverse host species.

In contrast, the host specialist *M. acridum* tended to prioritize C intake on early sampling days regardless of the P:C starting point, before switching to P on later sampling days (Fig. 4B, Table S3). This early-infection prioritization of C may correspond to a strategy of growth maximization during early infection stages[20,25], before switching to P used to induce sporulation at later stages.

## DISCUSSION

Our results support the framework modeling hosts as finite nutritional landscapes where nutritionally distinct tissues provide opportunities for niche partitioning by pathogens differing in developmental stage and life history. In so doing, we move beyond ‘food-level’ analyses of host specificity (e.g., locust specialist) and resolve the nutritional niche dimensions that govern pathogen performance.

Perhaps, it is not surprising that conventional labels of host specificity did not neatly predict fundamental nutritional niche (FNN) breadths of three *Metarhizium* species. Afterall, it is possible that the landscape of tissues with different P and C blends in a single locust host rivals the nutritional differences between landscapes found across hosts of different insect orders[1]. Yet, the niche-based framework we developed successfully resolved potentially general features of nutrient-specific growth and fitness whose niche dimensions can be compared across diverse pathogens, and across other types of species interactions from mutualists to interspecific competitors.

Our results also show the value of combining measures of pathogen FNN with measures of their nutritional foraging behavior. Specifically, host specificity did not mediate FNN breadths for three traits linked to fitness (growth area, timing and number of spores), but it did mediate nutritional intake dynamics. Specifically, the specialist (*M. acridum*) was more nutritionally selective than the generalists (*M. anisopliae*, *M. robertsii*) that tended to consume nutrients in the provided ratio.

Specifically, early-infection C-specific foraging by the locust-specialist *M. acridum* may be a life history adaptation to fast hyphal growth initially capitalizing on readily accessible C in hemolymph. Perhaps, these specialists can ‘anticipate’ that the early infection P deficit can eventually be redressed in the nutritionally predictable locust-host foraging landscape. Moreover, the switch to P-demanding spore production can occur more easily when these nutrients are liberated through nectrophic host decomposition.

In contrast, generalist *M. anisopliae* and *M. robertsii* appear to employ an opportunistic nutritional foraging strategy perhaps reflecting their uncertain future access to insect-derived nutrients. Specifically, unregulated P and C foraging may reflect that these generalists subsist asexually on nutrients derived from plant roots in the rhizosphere and from decaying organic matter in the soil, while only occasionally and opportunistically infecting insects[23]. More generally, we capitalized on the theoretical framework from community ecology to test whether a pathogen’s FNN dimensions are linked to their host specificity. We could then link changes in pathogen FNN dimensions to ontogenetic shifts from growth to proliferation to reproduction and dispersal. While we focus on nutrition, the niche of a pathogen is a composite of the availability of host nutrients and the capacity of the host immune response to minimize pathogen virulence[39,40].

It will be exciting to explore how host immune defenses mediate pathogen FNN dimensions in terms of proximate needs and host-specificity adaptations. For instance, no terrestrial ecosystem would be expected to actively inhibit potential invaders or dynamically screen among competitors. We propose that the nutritional niche framework is well suited to this task since compounds like host-derived anti-fungal peptides[41,42] can be added to nutritional media to resolve fine-scale interactions between P, C, and defensive compounds.

These interactions can be explored in three dimensional landscapes using right-angled mixture triangles[3] that can also substitute other nutrients hypothesized to govern immune performance like phosphorus [43,44] and iron[3,10,11]. In turn, this theoretical and empirical toolbox can be extended to other bacterial and fungal pathogens[24] and perhaps even to tumors[45] whose growth can be mediated by microscale nutritional environments within hosts.

## AUTHOR CONTRIBUTIONS

Conceptualization and Methodology, H.H.D.F.L. and J.Z.S.; Investigation, H.H.D.F.L, Z.S., P.J.D.N.N., E.B.L., A.K.K.H., and J.Z.S.; Writing – Original Draft, H.H.D.F.L. and J.Z.S.; Writing – Review and Editing, H.H.D.F.L. and J.Z.S.; Funding Acquisition, H.H.D.F.L. and J.Z.S.; Resources, H.H.D.F.L. and J.Z.S.

## METHODS

### Insect and fungal isolates

Adult *Locusta migratoria* were obtained from www.monis.dk and maintained in large groups of 50-100 individuals at room temperature (21±1 °C) with access to a heating lamp and fed every 2-3 days with fresh lettuce.

Three *Metarhizium* species (n = three isolates each of *M. acridum*, *M. anisopliae*, and two isolates of *M. robertsii*) maintained as glycerol stock solutions of conidia at -80°C at the Section for Organismal Biology, University of Copenhagen (Supplementary Table S4). Isolates were selected based on: 1) having similar growth and sporulation traits on standard media, and 2) targeting a geographically broad area of initial fungal collection. Fungal isolates were cultured on ¼ dilution of standard Sabouraud dextrose agar with yeast (SDAY/4: 2.5 g L-1 peptone, 10 g L-1 dextrose, 2.5 g L-1 yeast extract, 20 g L-1 agar) buffered a pH = 6,5 in constant darkness at 23°C. Each isolate was sub-cultured at most two times from being revived from the freezer-stock prior to inclusion in the study.

### Quantifying macronutrient contents of locust tissues

Adult female *Locusta migratoria* were obtained from www.monis.dk and maintained in large groups of 50-100 individuals at room temperature (21±1 °C) with access to a heating lamp and *ad lib* fresh lettuce replaced every two to three days. Live male and female locusts were weighed before making a small incision ventrally between the thorax and abdomen to extract hemolymph with a pipette. Locusts were euthanized by severing the head from the thorax. Dissections were carried out in insect physiological saline (IPS) buffer[46] by first carefully removing the entire digestive tract. The reproductive organs, fat body tissue, muscle tissue from the hind legs, and central nervous tissue from the brain were then collected in separate tubes and freeze dried.

To extract proteins (P) and carbohydrates (C), 2-µg freeze dried material from each tissue sample (nervous tissue from the brain had to be pooled from three to five individuals), was crushed with a pestle in 1-mL 0.1 M NaOH and centrifuged at 15.000 rpm for 15 min[47]. Protein content was measured using the Bradford colorimetric method by placing 2.5 µl of the supernatant with 250 µl of Bradford reagent (Sigma-Aldrich, B6916). Absorbance was read for these samples at 595 nm after an incubation period lasting 20 min. Readings of Bovine Serum Albumin standard at the same wavelength were used to generate a standard curve. Carbohydrate content was measured with a phenol-sulfuric acid method using the Total Carbohydrate Assay Kit (Sigma-Aldrich, MAK104) as described in the manual by the manufacturer. Carbohydrate content was determined with a spectrophotometer reading at 490 nm using a 2-mg/mL Glucose solution to generate a standard curve. Protein and carbohydrate contents were obtained by comparing optical density (OD) readings with the respective standard curves and are presented as mg/mL.

### Entomopathogenic fungal isolates

We studied three *Metarhizium* species (n = three isolates each of *M. acridum*, *M. anisopliae*, and two of *M. robertsii*) maintained long term in glycerol stock solutions of conidia at -80°C at the Section for Organismal Biology, University of Copenhagen (Supplementary Table S4). Isolates were selected based on: 1) having similar growth and sporulation traits on standard media, and 2) targeting a geographically broad area of initial fungal collection. Fungal isolates were cultured on ¼ dilution of standard Sabouraud dextrose agar with yeast (SDAY/4: 2.5-g L-1 peptone, 10-g L-1 dextrose, 2.5-g L-1 yeast extract, 20-g L-1 agar) buffered a pH = 6.5 in darkness at 23°C. Each isolate was sub-cultured at most two times after being revived from the freezer-stock prior to inclusion in the study.

### Fungus culturing and measurement of fungal growth

Conidia suspensions of the *Metarhizium* isolates were prepared by gently rubbing the surface of sporulating fungal cultures with a sterile borosilicate Drigalski spatula while adding 10-ml sterile 0,05% Triton X-100 (Merck) ddH_2_O solution. Conidium suspensions were washed to remove agar and fungal residues by centrifugation of the spore suspension at 3000 rpm for 3 min (Sigma 2-16KL). The supernatant was then discarded and this process was repeated. The cleaned conidia were then suspended in 10 ml of 0.05% Triton X-100 and this stock solution was serially diluted. We then used a 0.2-mm Fuchs-Rosenthal bright line hemocytometer to quantify spore concentrations by counting four, 16-cell squares under a microscope (Olympus BH-2).

Germination rates of all conidia suspensions were tested by spreading 100 µl of diluted conidia suspension on a SDAY/4 agar plate (100 × 15 mm) with a sterile Drigalski spatula. After a 24-h incubation at 23°C in darkness, we randomly selected two 100 x 100 mm squares of the inoculated medium and transferred them to a microscopy slide and assessed germination of 100 conidia in each square. The viability of conidia was evaluated on the same day as the suspension would be used for fungal inoculation, and only isolates showing > 90% germination rate were used. To obtain fungal growth rates, we point-inoculated Petri dishes (60 × 15-mm diameter) containing ca. 12.5 ml SDAY/4 media by adding 120 conidia (15 µl of a 8.000 conidia mL^-1^ conidial suspension) in the center. Following inoculation, Petri dishes were placed in a sterile bench for from 30 to 60 minutes allowing the spore droplet to soak into the solid agar-based media. Petri dishes were then sealed with parafilm and incubated in darkness at 23°C. The plates were monitored daily for any contamination and the area of the fungal colony measured every other day.

### Measuring the fundamental nutritional niche (FNN) of Metarhizium isolates

Nine *Metarhizium* isolates were inoculated on Petri dishes with one of 36 nutritionally defined diets spanning nine protein:carbohydrate (P:C) ratios (16:1, 8:1, 5:1, 3:1, 1:1, 1:3, 1:5, 1:8, 1:16 P:C) and four protein + carbohydrate (P + C) concentrations (4, 8, 20, and 50 g/L P + C). Diets were prepared by combining 1.6 w/v % of microbiological agar (Sigma-Aldrich), soluble starch (Sigma-Aldrich), sucrose (Sigma-Aldrich), Bacto peptone (Becton Dickinson, BD), Bacto tryptone (BD), Trypticase Peptone (BD), and Vanderzant vitamin mixture (Sigma) at 2% total dry mass. Diet recipes were adapted from the study of Shik et al.[48] and are provided in (Supplementary Table S5). Dry diet ingredients were weighed to the nearest 0.001 g on a scale (Denver Instrument, SI-234), suspended in 500 ml distilled water, stirred for five to ten minutes while the pH was adjusted to 6.5, and then autoclaved at 121°C. If necessary (for higher diet concentrations), we slowly mixed the autoclaved media again using a stirring magnet. Each Petri dish (60×15 mm diameter) contained approximately 12.5 ml of media and was stored at room temperature for no longer than one week before being used.

We point-inoculated each Petri dish (60 × 15 mm diameter) by adding 120 conidia (5 µl of a 24000 conidia mL^-1^ conidial suspension) in the center. Following inoculation, Petri dishes were placed in a sterile bench for 30-60 minutes allowing the spore droplet to soak into the solid agar-based media. Petri dishes were then sealed with parafilm and then incubated in darkness at 23°C. The plates were monitored daily for contamination. Each of the 36 diet treatments was replicated three times for each of the nine fungal isolates (N = 972 Petri dishes). After 11 days of growth, each Petri dish was photographed with a camera (Canon EOS 700) in a photo box ensuring uniform light and focus area and equipped with a reference ruler.

Mycelial growth area (mm^2^) was obtained by analyzing photographs using ImageJ (NIH; v1.52a). Conidia number was obtained from each Petri dish after 15 days of incubation by adding 3 ml of sterile Triton-X 0,05% solution and then carefully scraping the surface with a sterile spatula to suspend the spores in the solution. We transferred each suspension to a 15 ml Falcon tube and froze it at -20°C. After defrosting each sample, we performed serial dilutions to adjust each spore concentration into a measurable range. We counted 20 µl of diluted spore suspension in a 0.2-mm Fuchs-Rosenthal bright line cytometer with two or three technical replicates of each isolate and nutritional diet combination.

To estimate the effects of P:C diet on the onset of sporulation, we monitored each Petri dish daily for 11 days and used a modified scoring scheme from Fernandes et al.[49] where the fungus color was qualitatively evaluated on an eight-level scale indicating developmental stage from white (hyphal growth) to dark green (mature sporulation). We detail this color scoring scheme in Supplementary Table S6 and provide representative images of each species at each color level in Figure S2. Most isolates exhibited some green coloration by Day 5, but some diet treatments never sporulated by Day 11. For subsequent analyses we analyzed the onset of sporulation in units of days before the end of the experiment at Day 11 and focused on the first sign of green spores (Category 5, or 6).

We visualized FNN heatmaps using the fields package[50] in R (4.3.1)[51], plotting the response variables growth area (mm^2^), log_10_(spore number), and onset day of sporulation across the 36 P:C diet treatments. Red areas indicate high values of response variables and blue areas indicate low values. We set the topological resolution of response surfaces to λ=0.0005 as a smoothing factor[48] and generated performance isoclines using non-parametric thin-plate splines. We used least-square regressions to assess the significance of the linear and quadratic terms (and their linear interaction) for each dependent variable across the P and C diet treatments. These analyses were performed on the mean values across isolates for *M. anisopliae* (n = 3), *M. acridum* (n = 3), and *M. robertsii* (n = 2) used to generate the composite species-level figures (Table S1), and at the level of each isolate (Table S2).

### Measuring Metarhizium nutrient depletion of nutritionally defined media

We next used an *in vitro* liquid media approach to test for nutrient-specific foraging of developing *Metarhizium* fungi. We prepared four nutritionally defined media treatments in a factorial design (P:C: 1:3, 3:1, P+C: 15 g/L, 50 g/L) using the approach previously described, but this time without agar. Petri dishes (90-mm diameter) were first filled with a single layer of sterile 5-mm glass beads and then filled with 10 mL of liquid media to cover the glass beads. We focused on one isolate of each species (*M. anisopliae* ARSEF_549, *M. acridum* ARSEF_324, and *M. robertsii* KVL_12-35; Table S4), with ten Petri dish replicates per isolate and media combination. Each Petri dish was inoculated with a 5 x 5-mm agar plug (SDAY/4 media) containing fungal hyphae placed upside down in the center of the dish. Each Petri dish was then sealed with parafilm. Every 48 hours, we gently shook each Petri dish, before opening inside a sterile hood and sampling 200 µl of the liquid growth media with a sterile pipette. Samples were stored frozen at -20°C until analysis.

We measured carbohydrate content using the phenol-sulfuric acid method with the Total Carbohydrate Assay Kit (Cell Biolabs, STA-682) as described above. For the protein measurements, we used 2 µl of liquid media diluted 15 times in phosphate buffered saline (PBS) solution (pH = 7.4) in a micro-volume spectrophotometer at 280 nm. Equal amounts (1:1:1) of bacto peptone, bacto tryptone, trypticase peptone suspended in PBS buffer were used as standard. In total, we sampled media for protein and carbohydrate on day 0 (freshly made media), day 2 (48 hours), day 4 (96 hours), and day 6 (144 hours) after initial inoculation.

### Quantification and statistical analyses

All statistical analyses were performed using R Studio (4.3.1)[51]. Details of analyses are given in the sections *Measuring the fundamental nutritional niche (FNN) of Metarhizium isolates* and *Measuring Metarhizium nutrient depletion of nutritionally defined media*.

## Supporting information

ZIP-file containing data and code

