## Supplementary tables and figures for "Insect hosts are nutritional landscapes navigated by fungal pathogens"

### Supplementary material

#### Supplementary tables

**Suppl. table S1**

**Supplementary table S1. Regression analyses supporting heatmaps of mean species cultivar growth area ($mm^2$), color category (green color representing onset of sporulation), and spore number ($log_{10}$ spore number).** Presented are least square regression significances for both linear and quadratic terms for protein (P) and carbohydrate (C) composition in agar plate substrates, as well as the P x C interaction. Significant results for univariate models support the overall interpretation of FNN heatmaps visualizing fungal growth area ($mm^2$) across the 36 protein and carbohydrate combinations. The across-isolate (species mean) analyses are based on mean-value analysis, averaging across the P:C treatment means for each isolate, with three isolates for *M. anisopliae* ESALQ_1116, ESALQ_1604, ARSEF_549), three isolates for *M. acridum* (ARSEF_7486, ARSEF_3609, ARSEF_324), and two isolates for *M. robertsii* (KVL_12-35, KVL_13-12). The same analyses were also performed at the individual isolate level (Table S5). See Methods, Table S2 and S3, and Figure S3 for more information about how growth area, color category, and spore number were calculated.

| **Significance test for univariate models** | | | | | |
|---|---|---|---|---|---|
| **Model** | **DF model** | **DF error** | **$R^2$ adjusted** | **F** | ***P*** |
| *M. anisopliae* | | | | | |
| Area | 5 | 30 | 0.87 | 47.04 | **< 0.0001** |
| Color category | 5 | 30 | 0.79 | 26.69 | **< 0.0001** |
| Spore number | 5 | 30 | 0.82 | 32.96 | **< 0.0001** |
| *M. acridum* | | | | | |
| Area | 5 | 30 | 0.78 | 25.23 | **< 0.0001** |
| Color | 5 | 30 | 0.79 | 27.79 | **< 0.0001** |
| Spore number | 5 | 30 | 0.44 | 6.46 | **0.0003** |
| *M. robertsii* | | | | | |
| Area | 5 | 30 | 0.81 | 31.37 | **0.0001** |
| Color | 5 | 30 | 0.41 | 5.93 | **0.0006** |

| | | | | | |
|---|---|---|---|---|---|
| Spore | 5 | 30 | 0.39 | 5.51 | **0.001** |

| **Univariate tests of Parameter estimates** | | | | |
|---|---|---|---|---|
| **Isolate** | **Parameter** | **DF** | **F** | **P** |
| *M. anisopliae* | | | | |
| Area | P | 1 | 117.84 | **< 0.0001** |
| | C | 1 | 0.55 | 0.464 |
| | $P^2$ | 1 | 66.30 | **< 0.0001** |
| | $C^2$ | 1 | 2.58 | 0.119 |
| | PC | 1 | 7.02 | **0.013** |
| | Error | 30 | | |
| Color | P | 1 | 33.35 | **< 0.0001** |
| | C | 1 | 15.28 | **0.0005** |
| | $P^2$ | 1 | 9.48 | **0.004** |
| | $C^2$ | 1 | 7.04 | **0.013** |
| | PC | 1 | 0.05 | 0.821 |
| | Error | 30 | | |
| Spore number | P | 1 | 10.72 | **0.001** |
| | C | 1 | 23.95 | **< 0.0001** |
| | $P^2$ | 1 | 0.44 | 0.741 |
| | $C^2$ | 1 | 0.07 | 0.603 |
| | PC | 1 | 20.71 | **< 0.0001** |
| | Error | 30 | | |
| *M. acridum* | | | | |
| Area | P | 1 | 85.16 | **< 0.0001** |
| | C | 1 | 0.78 | 0.384 |
| | $P^2$ | 1 | 61.53 | **< 0.0001** |
| | $C^2$ | 1 | 2.84 | 0.102 |
| | PC | 1 | 6.57 | **0.016** |
| | Error | 30 | | |
| Color | P | 1 | 23.06 | **< 0.0001** |

| | | | | |
|---|---|---|---|---|
| | C | 1 | 5.78 | **0.023** |
| | $P^2$ | 1 | 14.89 | **0.0006** |
| | $C^2$ | 1 | 6.80 | **0.014** |
| | PC | 1 | 17.74 | **0.0002** |
| | Error | 30 | | |
| Spore number | P | 1 | 0.98 | 0.329 |
| | C | 1 | 14.43 | **0.0007** |
| | $P^2$ | 1 | 0.09 | 0.769 |
| | $C^2$ | 1 | 4.00 | 0.055 |
| | PC | 1 | 1.63 | 0.212 |
| | Error | 30 | | |
| *M. robertsii* | | | | |
| Area | P | 1 | 124.21 | **< 0.0001** |
| | C | 1 | 3.87 | 0.058 |
| | $P^2$ | 1 | 77.87 | **< 0.0001** |
| | $C^2$ | 1 | 0.43 | 0.516 |
| | PC | 1 | 0.07 | 0.792 |
| | Error | 30 | | |
| Color | P | 1 | 4.30 | **0.047** |
| | C | 1 | 3.37 | 0.076 |
| | $P^2$ | 1 | 2.23 | 0.145 |
| | $C^2$ | 1 | 2.31 | 0.139 |
| | PC | 1 | 1.78 | 0.192 |
| | Error | 30 | | |
| Spore number | P | 1 | 7.38 | **0.011** |
| | C | 1 | 13.16 | **0.001** |
| | $P^2$ | 1 | 3.39 | 0.076 |
| | $C^2$ | 1 | 5.74 | **0.023** |
| | PC | 1 | 1.37 | 0.251 |
| | Error | 30 | | |

| Parameter estimates | | | |
|---|---|---|---|
| Isolate | Parameter | Estimate | **P** |
| *M. anisopliae* | | | |
| Area | P | 0.40 | **< 0.0001** |
| | C | 0.08 | **0.038** |
| | $P^2$ | -0.01 | **< 0.0001** |
| | $C^2$ | -0.00 | 0.119 |
| | PC | -0.00 | **0.013** |
| Color | P | -0.33 | **< 0.0001** |
| | C | -0.22 | **0.005** |
| | $P^2$ | 0.00 | **0.004** |
| | $C^2$ | 0.00 | **0.013** |
| | PC | -0.00 | 0.820 |
| Spore number | P | 0.08 | 0.100 |
| | C | 0.02 | 0.697 |
| | $P^2$ | -0.00 | 0.513 |
| | $C^2$ | -0.00 | 0.787 |
| | PC | -0.01 | **< 0.0001** |
| *M. acridum* | | | |
| Area | P | 0.16 | **< 0.0001** |
| | C | 0.04 | **0.033** |
| | $P^2$ | -0.00 | **< 0.0001** |
| | $C^2$ | -0.00 | 0.102 |
| | PC | -0.00 | **0.016** |
| Color | P | -0.46 | **< 0.0001** |
| | C | -0.32 | **0.0001** |
| | $P^2$ | 0.01 | **0.0006** |
| | $C^2$ | 0.00 | **0.014** |
| | PC | 0.01 | **0.0002** |
| Spore number | P | -0.00 | 0.967 |
| | C | -0.18 | **0.028** |

| | | | |
|---|---|---|---|
| | $P^2$ | 0.00 | 0.770 |
| | $C^2$ | 0.00 | 0.055 |
| | PC | -0.00 | 0.212 |
| *M. robertsii* | | | |
| Area | P | 0.49 | **< 0.0001** |
| | C | -0.08 | 0.164 |
| | $P^2$ | -0.01 | **< 0.0001** |
| | $C^2$ | 0.00 | 0.516 |
| | PC | -0.00 | 0.792 |
| Color | P | -0.24 | **0.020** |
| | C | -0.22 | **0.031** |
| | $P^2$ | 0.00 | 0.146 |
| | $C^2$ | 0.00 | 0.139 |
| | PC | 0.01 | 0.192 |
| Spore number | P | -0.18 | 0.146 |
| | C | -0.27 | **0.034** |
| | $P^2$ | 0.00 | 0.076 |
| | $C^2$ | 0.01 | **0.023** |
| | PC | -0.01 | 0.251 |

**Suppl. table S2**

**Supplementary table S2 Regression analyses supporting heatmaps of individual isolate cultivar growth area ($mm^2$), color category, and spore number ($log_{10}$ spore number).** Presented are least square regression significances for both linear and quadratic terms for protein (P) and carbohydrate (C) composition in agar plate substrates, as well as the P x C interaction. Significant results for univariate models support the overall interpretation of FNN heatmaps visualizing fungal growth area ($mm^2$) across the 36 protein and carbohydrate combinations. We provide separate analyses of each of the 8 isolates (*M. anisopliae* (ESALQ_1116, ESALQ_1604, ARSEF_549), *M. acridum* (ARSEF_7486, ARSEF_3609, ARSEF_324), *M. robertsii* (KVL_12-35, KVL_13-12). See Methods, Table S2 and S3, and Figure S3 for more information about how growth area, color, and spore number were calculated.

| **Significance test for univariate models** | | | | | |
|---|---|---|---|---|---|
| **Model** | **DF model** | **DF error** | **$R^2$ adjusted** | **F** | ***P*** |
| *M. anisopliae* [ESALQ_1116] | | | | | |
| Area | 5 | 30 | 0.84 | 37.81 | **< 0.0001** |
| Color | 5 | 30 | 0.79 | 27.45 | **< 0.0001** |
| Spore number | 5 | 30 | 0.76 | 22.58 | **< 0.0001** |
| *M. anisopliae* [ESALQ_1604] | | | | | |
| Area | 5 | 30 | 0.59 | 10.98 | **< 0.0001** |
| Color | 5 | 30 | 0.59 | 10.87 | **< 0.0001** |
| Spore number | 5 | 30 | 0.07 | 1.56 | 0.201 |
| *M. anisopliae* [ARSEF_549] | | | | | |
| Area | 5 | 30 | 0.79 | 27.74 | **< 0.0001** |
| Color | 5 | 30 | 0.91 | 76.19 | **< 0.0001** |
| Spore number | 5 | 30 | 0.81 | 30.29 | **< 0.0001** |
| *M. acridum* [ARSEF_7486] | | | | | |
| Area | 5 | 30 | 0.75 | 21.61 | **< 0.0001** |
| Color | 5 | 30 | 0.74 | 20.92 | **< 0.0001** |
| Spore number | 5 | 30 | 0.26 | 3.42 | **< 0.0001** |
| *M. acridum* [ARSEF_3609] | | | | | |
| Area | 5 | 30 | 0.88 | 54.34 | **< 0.0001** |
| Color | 5 | 30 | 0.53 | 8.90 | **< 0.0001** |

| | | | | | |
|---|---|---|---|---|---|
| Spore number | 5 | 30 | 0.55 | 9.61 | **< 0.0001** |
| *M. acridum* [ARSEF_324] | | | | | |
| Area | 5 | 30 | 0.38 | 5.23 | **0.001** |
| Color | 5 | 30 | 0.84 | 37.55 | **< 0.0001** |
| Spore number | 5 | 30 | 0.11 | 1.85 | 0.134 |
| *M. robertsii* [KVL_12-35] | | | | | |
| Area | 5 | 30 | 0.47 | 7.24 | **0.0001** |
| Color | 5 | 30 | 0.25 | 3.34 | **0.016** |
| Spore | 5 | 30 | 0.56 | 9.87 | **< 0.0001** |
| *M. robertsii* [KVL_13-12] | | | | | |
| Area | 5 | 30 | 0.88 | 51.96 | **< 0.0001** |
| Color | 5 | 30 | 0.47 | 7.18 | **0.0002** |
| Spore number | 5 | 30 | 0.21 | 2.81 | **0.033** |

| | | | | |
|---|---|---|---|---|
| **Univariate tests of Parameter estimates** | | | | |
| **Isolate** | **Parameter** | **DF** | **F** | **P** |
| *M. anisopliae* [ESALQ_1116] | | | | |
| Area | P | 1 | 83.34 | **< 0.0001** |
| | C | 1 | 0.28 | 0.599 |
| | $P^2$ | 1 | 45.99 | **< 0.0001** |
| | $C^2$ | 1 | 1.08 | 0.308 |
| | PC | 1 | 7.95 | **0.008** |
| | Error | 30 | | |
| Color | P | 1 | 64.95 | **< 0.0001** |
| | C | 1 | 4.62 | **0.040** |
| | $P^2$ | 1 | 31.26 | **< 0.0001** |
| | $C^2$ | 1 | 3.91 | 0.057 |
| | PC | 1 | 0.61 | 0.439 |
| | Error | 30 | | |
| Spore number | P | 1 | 12.67 | **0.001** |
| | C | 1 | 21.25 | **< 0.0001** |

| | | | | |
|---|---|---|---|---|
| | $P^2$ | 1 | 0.11 | 0.741 |
| | $C^2$ | 1 | 0.28 | 0.603 |
| | PC | 1 | 59.20 | **< 0.0001** |
| | Error | 30 | | |
| *M. anisopliae* [ESALQ_1604] | | | | |
| Area | P | 1 | 29.29 | **< 0.0001** |
| | C | 1 | 3.69 | 0.064 |
| | $P^2$ | 1 | 14.01 | **0.0008** |
| | $C^2$ | 1 | 1.41 | 0.245 |
| | PC | 1 | 0.08 | 0.786 |
| | Error | 30 | | |
| Color | P | 1 | 7.20 | **0.012** |
| | C | 1 | 13.16 | **0.001** |
| | $P^2$ | 1 | 1.63 | 0.212 |
| | $C^2$ | 1 | 7.046 | **0.013** |
| | PC | 1 | 0.015 | 0.903 |
| | Error | 30 | | |
| Spore number | P | 1 | 0.01 | 0.916 |
| | C | 1 | 0.63 | 0.433 |
| | $P^2$ | 1 | 0.07 | 0.79 |
| | $C^2$ | 1 | 0.12 | 0.732 |
| | PC | 1 | 0.490 | 0.489 |
| | Error | 30 | | |
| *M. anisopliae* [ARSEF_549] | | | | |
| Area | P | 1 | 57.79 | **< 0.0001** |
| | C | 1 | 10.63 | **< 0.0001** |
| | $P^2$ | 1 | 36.17 | **0.003** |
| | $C^2$ | 1 | 9.03 | **0.005** |
| | PC | 1 | 5.30 | **0.028** |
| | Error | 30 | | |
| Color | P | 1 | 43.41 | **< 0.0001** |

| | | | | |
|---|---|---|---|---|
| | C | 1 | 25.29 | **< 0.0001** |
| | $P^2$ | 1 | 3.07 | 0.090 |
| | $C^2$ | 1 | 2.10 | 0.157 |
| | PC | 1 | 2.65 | 0.114 |
| | Error | 30 | | |
| Spore number | P | 1 | 8.03 | **0.008** |
| | C | 1 | 12.95 | **0.001** |
| | $P^2$ | 1 | 1.46 | 0.237 |
| | $C^2$ | 1 | 1.84 | **0.186** |
| | PC | 1 | 50.86 | **< 0.0001** |
| | Error | 30 | | |
| *M. acridum* [ARSEF_7486] | | | | |
| Area | P | 1 | 88.40 | **< 0.0001** |
| | C | 1 | 2.63 | 0.116 |
| | $P^2$ | 1 | 55.63 | **< 0.0001** |
| | $C^2$ | 1 | 0.54 | 0.466 |
| | PC | 1 | 0.08 | 0.778 |
| | Error | 30 | | |
| Color | P | 1 | 29.05 | **< 0.0001** |
| | C | 1 | 4.55 | **0.041** |
| | $P^2$ | 1 | 19.10 | **0.0001** |
| | $C^2$ | 1 | 6.69 | **0.015** |
| | PC | 1 | 11.44 | **0.002** |
| | Error | 30 | | |
| Spore number | P | 1 | 0.060 | 0.809 |
| | C | 1 | 7.68 | **0.009** |
| | $P^2$ | 1 | 0.85 | 0.365 |
| | $C^2$ | 1 | 1.24 | 0.275 |
| | PC | 1 | 4.98 | **0.033** |
| | Error | 30 | | |

| | | | | |
|---|---|---|---|---|
| *M. acridum* [ARSEF_3609] | | | | |
| Area | P | 1 | 97.99 | **< 0.0001** |
| | C | 1 | 15.19 | **0.0005** |
| | $P^2$ | 1 | 44.61 | **< 0.0001** |
| | $C^2$ | 1 | 14.10 | **0.0007** |
| | PC | 1 | 3.54 | **0.070** |
| | Error | 30 | | |
| Color | P | 1 | 19.90 | **0.018** |
| | C | 1 | 0.090 | 0.868 |
| | $P^2$ | 1 | 8.890 | 0.106 |
| | $C^2$ | 1 | 0.153 | 0.828 |
| | PC | 1 | 15.26 | **0.037** |
| | Error | 30 | | |
| Spore number | P | 1 | 5.10 | **0.031** |
| | C | 1 | 10.93 | **0.002** |
| | $P^2$ | 1 | 1.76 | 0.195 |
| | $C^2$ | 1 | 0.90 | 0.349 |
| | PC | 1 | 4.64 | **0.039** |
| | Error | 30 | | |
| *M. acridum* [ARSEF_324] | | | | |
| Area | P | 1 | 16.14 | **0.0004** |
| | C | 1 | 0.24 | 0.626 |
| | $P^2$ | 1 | 18.76 | **0.0002** |
| | $C^2$ | 1 | 1.42 | 0.243 |
| | PC | 1 | 6.63 | **0.015** |
| | Error | 30 | | |
| Color | P | 1 | 17.01 | **0.0003** |
| | C | 1 | 21.71 | **< 0.0001** |
| | $P^2$ | 1 | 14.05 | **0.0008** |
| | $C^2$ | 1 | 17.63 | **0.0002** |
| | PC | 1 | 25.92 | **< 0.0001** |

| | | | | |
|---|---|---|---|---|
| | Error | 30 | | |
| Spore number | P | 1 | 0.09 | 0.765 |
| | C | 1 | 6.07 | **0.020** |
| | $P^2$ | 1 | 0.63 | 0.630 |
| | $C^2$ | 1 | 0.06 | 0.062 |
| | PC | 1 | 0.91 | 0.912 |
| | Error | 30 | | |
| *M. robertsii* [KVL_12-35] | | | | |
| Area | P | 1 | 28.68 | **< 0.0001** |
| | C | 1 | 0.01 | 0.927 |
| | $P^2$ | 1 | 16.96 | **0.0003** |
| | $C^2$ | 1 | 0.05 | 0.824 |
| | PC | 1 | 1.35 | 0.254 |
| | Error | 30 | | |
| Color | P | 1 | 1.36 | 0.253 |
| | C | 1 | 1.48 | 0.233 |
| | $P^2$ | 1 | 0.05 | 0.833 |
| | $C^2$ | 1 | 0.25 | 0.620 |
| | PC | 1 | 0.05 | 0.832 |
| | Error | 30 | | |
| Spore number | P | 1 | 6.06 | **0.020** |
| | C | 1 | 18.70 | **0.0002** |
| | $P^2$ | 1 | 1.08 | 0.304 |
| | $C^2$ | 1 | 3.72 | 0.063 |
| | PC | 1 | 7.61 | **0.010** |
| | Error | 30 | | |
| *M. robertsii* [KVL_13-12] | | | | |
| Area | P | 1 | 183.67 | **< 0.0001** |
| | C | 1 | 12.87 | **0.001** |
| | $P^2$ | 1 | 118.86 | **< 0.0001** |

| | | | | |
|---|---|---|---|---|
| | $C^2$ | 1 | 2.42 | 0.130 |
| | PC | 1 | 4.55 | **0.041** |
| | Error | 30 | | |
| Color | P | 1 | 6.50 | **0.016** |
| | C | 1 | 4.21 | **0.049** |
| | $P^2$ | 1 | 6.40 | **0.017** |
| | $C^2$ | 1 | 5.10 | **0.031** |
| | PC | 1 | 7.31 | **0.011** |
| | Error | 30 | | |
| Spore number | P | 1 | 6.41 | **0.017** |
| | C | 1 | 6.07 | **0.020** |
| | $P^2$ | 1 | 5.15 | **0.031** |
| | $C^2$ | 1 | 6.00 | **0.020** |
| | PC | 1 | 0.l7 | 0.682 |
| | Error | 30 | | |

| **Parameter estimates** | | | |
|---|---|---|---|
| **Isolate** | **Parameter** | **Estimate** | **P** |
| *M. anisopliae* [ESALQ_1116] | | | |
| Area | P | 0.44 | **< 0.0001** |
| | C | -0.01 | **< 0.0001** |
| | $P^2$ | 0.06 | 0.223 |
| | $C^2$ | -0.00 | 0.308 |
| | PC | -0.01 | **0.008** |
| Color | P | -0.55 | **< 0.0001** |
| | C | -0.17 | **0.036** |
| | $P^2$ | 0.01 | **< 0.0001** |
| | $C^2$ | 0.00 | 0.057 |
| | PC | 0.00 | 0.439 |
| Spore number | P | 0.12 | 0.100 |
| | C | 0.06 | 0.402 |
| | $P^2$ | -0.00 | 0.741 |

| | | | |
|---|---|---|---|
| | $C^2$ | 0.00 | 0.603 |
| | PC | -0.02 | **< 0.0001** |
| *M. anisopliae* [ESALQ_1604] | | | |
| Area | P | 0.25 | **< 0.0001** |
| | C | -0.08 | 0.177 |
| | $P^2$ | -0.00 | 0.001 |
| | $C^2$ | 0.00 | 0.245 |
| | PC | -0.00 | 0.786 |
| Color | P | -0.25 | **0.033** |
| | C | -0.33 | **0.005** |
| | $P^2$ | 0.00 | 0.211 |
| | $C^2$ | 0.01 | 0.013 |
| | PC | 0.00 | 0.903 |
| Spore number | P | 0.12 | 0.100 |
| | | 0.06 | 0.402 |
| | $P^2$ | -0.00 | 0.741 |
| | $C^2$ | 0.00 | 0.603 |
| | PC | -0.02 | **< 0.0001** |
| *M. anisopliae* [ARSEF_549] | | | |
| Area | P | 0.50 | **< 0.0001** |
| | C | 0.27 | **0.0004** |
| | $P^2$ | -0.01 | **< 0.0001** |
| | $C^2$ | 0.00 | **0.005** |
| | PC | -0.01 | **0.028** |
| Color | P | 0.05 | **0.0001** |
| | C | 0.05 | **0.004** |
| | $P^2$ | 0.00 | 0.090 |
| | $C^2$ | 0.00 | 0.157 |
| | PC | 0.00 | 0.114 |
| Spore number | P | 0.17 | 0.061 |
| | C | 0.12 | 0.193 |

| | | | |
|---|---|---|---|
| | $P^2$ | -0.00 | 0.237 |
| | $C^2$ | -0.00 | 0.186 |
| | PC | -0.02 | **< 0.0001** |
| *M. acridum* [ARSEF_7486] | | | |
| Area | P | 0.15 | **< 0.0001** |
| | C | -0.02 | 0.265 |
| | $P^2$ | -0.00 | **< 0.0001** |
| | $C^2$ | 0.00 | 0.466 |
| | PC | -0.00 | 0.778 |
| Color | P | -0.53 | **0.0001** |
| | C | -0.31 | **0.0008** |
| | $P^2$ | 0.01 | **0.0001** |
| | $C^2$ | 0.00 | **0.015** |
| | PC | 0.01 | **0.002** |
| Spore number | P | 0.03 | 0.139 |
| | C | -0.02 | 0.371 |
| | $P^2$ | -0.00 | 0.365 |
| | $C^2$ | 0.00 | 0.275 |
| | PC | -0.00 | **0.033** |
| *M. acridum* [ARSEF_3609] | | | |
| Area | P | 0.14 | **< 0.0001** |
| | C | 0.07 | **0.0002** |
| | $P^2$ | -0.00 | **< 0.0001** |
| | $C^2$ | -0.00 | **0.0007** |
| | PC | -0.00 | 0.070 |
| Color | P | -0.36 | **0.002** |
| | C | -0.12 | 0.256 |
| | $P^2$ | 0.00 | 0.106 |
| | $C^2$ | 0.00 | 0.828 |
| | PC | 0.01 | **0.037** |
| Spore number | P | 0.03 | 0.139 |

| | | | |
|---|---|---|---|
| | C | -0.02 | 0.371 |
| | | -0.00 | 0.365 |
| | $C^2$ | 0.00 | 0.275 |
| | PC | -0.00 | **0.033** |
| *M. acridum* [ARSEF_324] | | | |
| Area | P | 0.19 | **< 0.0001** |
| | C | 0.08 | **< 0.0001** |
| | $P^2$ | -0.00 | 0.064 |
| | $C^2$ | -0.00 | **0.0002** |
| | PC | -0.00 | **0.015** |
| Color | P | -0.36 | **0.002** |
| | C | -0.12 | 0.256 |
| | $P^2$ | 0.00 | 0.106 |
| | $C^2$ | 0.00 | 0.828 |
| | PC | 0.01 | **0.037** |
| Spore number | P | 0.03 | 0.861 |
| | C | -0.34 | **0.049** |
| | $P^2$ | -0.00 | 0.630 |
| | $C^2$ | 0.01 | 0.062 |
| | PC | 0.00 | 0.912 |
| *M. robertsii* [KVL_12-35] | | | |
| Area | P | 0.29 | **0.001** |
| | C | -0.06 | 0.451 |
| | $P^2$ | -0.01 | **0.0003** |
| | $C^2$ | -0.00 | 0.824 |
| | PC | 0.00 | 0.254 |
| Color | P | -0.10 | 0.424 |
| | C | -0.10 | 0.400 |
| | $P^2$ | 0.00 | 0.833 |
| | $C^2$ | 0.00 | 0.620 |
| | PC | -0.00 | 0.832 |

| | | | |
|---|---|---|---|
| Spore number | P | -0.04 | 0.731 |
| | C | -0.23 | 0.074 |
| | $P^2$ | 0.00 | 0.304 |
| | $C^2$ | 0.00 | 0.063 |
| | PC | -0.01 | **0.010** |
| *M. robertsii* [KVL_13-12] | | | |
| Area | P | 0.69 | **< 0.0001** |
| | C | -0.09 | 0.4115 |
| | $P^2$ | -0.01 | **< 0.0001** |
| | $C^2$ | 0.00 | 0.130 |
| | PC | -0.00 | **0.041** |
| Color | P | -0.39 | **0.001** |
| | C | -0.35 | **0.003** |
| | $P^2$ | 0.01 | **0.017** |
| | $C^2$ | 0.00 | **0.031** |
| | PC | 0.01 | **0.011** |
| Spore number | P | -0.04 | 0.731 |
| | C | -0.23 | 0.074 |
| | $P^2$ | 0.00 | 0.304 |
| | $C^2$ | 0.00 | 0.063 |
| | PC | -0.01 | **0.010** |

**Suppl. table S3**

**Supplementary table S3.** Linear regression analysis comparing fungal nutritional consumption in relation to the ratio of protein and carbohydrate in the media. The results of comparing the difference between coefficients of the linear model of *lm*(carbohydrates~protein) with the $H_0$ of a linear consumption of media ratios (either 3.00 or 0.33). Tests were conducted using the function *LinearHypothesis* from the *car* package (Fox & Weisberg, 2019) in R version 4.3.1.

| Media composition | Slope estimate | $H_0$ | F | *p* |
|---|---|---|---|---|
| ***M. anisopliae*** | | | | |
| 15 g/L, P:C = 1:3 | 2.15 | 3.00 | 2.3940 | 0.1528 |
| 15 g/L, P:C = 3:1 | 0.11 | 0.33 | **6.3541** | **0.0304** |
| 50 g/L, P:C = 1:3 | 1.54 | 3.00 | 2.9302 | 0.1177 |
| 50 g/L, P:C = 3:1 | 0.37 | 0.33 | 0.2295 | 0.6433 |
| ***M. robertsii*** | | | | |
| 15 g/L, P:C = 1:3 | 0.30 | 3.00 | **29.982** | **0.0003** |
| 15 g/L, P:C = 3:1 | 0.22 | 0.33 | 3.3850 | 0.0956 |
| 50 g/L, P:C = 1:3 | 0.84 | 3.00 | **19.646** | **0.0013** |
| 50 g/L, P:C = 3:1 | 0.40 | 0.33 | 0.3192 | 0.5846 |
| ***M. acridum*** | | | | |
| 15 g/L, P:C = 1:3 | 2.02 | 3.00 | 3.4500 | 0.0929 |
| 15 g/L, P:C = 3:1 | 0.20 | 0.33 | 1.1641 | 0.3060 |
| 50 g/L, P:C = 1:3 | 1.72 | 3.00 | 3.5239 | 0.0899 |
| 50 g/L, P:C = 3:1 | 0.36 | 0.33 | 0.0282 | 0.8700 |

*Fox, J. and Weisberg, S. (2019) An R Companion to Applied Regression, Third Edition, Sage.*

**Suppl. table S4**

**Supplementary table S4.** Isolates of *Metarhizium* used in current study. Isolates in bold were included in the nutritional consumption experiment.

| Species | Code | Isolate | Source | Country | Year | Notes |
|---|---|---|---|---|---|---|
| ***M. anisopliae*** | **A1** | **ARSEF_549** | **n.a.** | **Brazil** | **1980** | |
| *M. anisopliae* | A | ESALQ_1116 | *Scarabaeidae* | Brazil | 1993 | |
| *M. anisopliae* | B | ESALQ_1604 | n.a. | Brazil | n.a. | Biotech G, Biotech® Controle Biológico (Isolated from commercial isolate) |
| ***M. acridum*** | **B1** | **ARSEF_324** | ***Austracris guttulosa*** | **Australia** | **1979** | |
| *M. acridum* | C | ARSEF_7486 | *Ornithacris cavroisi* | Niger | 1992 | Active ingredient in commercial product *Green Muscle*™ |
| *M. acridum* | D | ARSEF_3609 | *Patanga succincta* | Thailand | 1992 | |
| ***M. robertsii*** | **E** | **KVL_12-35** | **Soil** | **Denmark** | **2012** | |
| *M. robertsii* | F | KVL_13-12 | Soil | Denmark | 2013 | |

**Suppl. Table S5**

**Supplementary table S5.** Agar media recipes for 36 fungal diets. All quantities are in grams (g) and total volume of media is 500 mL (Modified from Shik et al. 2016).

| P:C | Bacto Peptone | Trypticase Peptone | Bacto Tryptone | Sucrose | Starch | Agar | Vitamin |
|---|---|---|---|---|---|---|---|
| **Media concentration 4 g/l** | | | | | | | |
| 16:1 | 0.6676 | 0.6680 | 0.6759 | 0.0539 | 0.0539 | 8 | 0.4 |
| 8:1 | 0.6305 | 0.6309 | 0.6383 | 0.1065 | 0.1065 | 8 | 0.4 |
| 5:1 | 0.5911 | 0.5915 | 0.5984 | 0.1623 | 0.1623 | 8 | 0.4 |
| 3:1 | 0.5320 | 0.5323 | 0.5385 | 0.2460 | 0.2460 | 8 | 0.4 |
| 1:1 | 0.3548 | 0.3550 | 0.3590 | 0.4975 | 0.4975 | 8 | 0.4 |
| 1:3 | 0.1773 | 0.1775 | 0.1795 | 0.7488 | 0.7488 | 8 | 0.4 |
| 1:5 | 0.1182 | 0.1183 | 0.1197 | 0.8325 | 0.8325 | 8 | 0.4 |
| 1:8 | 0.0788 | 0.0789 | 0.0798 | 0.8883 | 0.8883 | 8 | 0.4 |
| 1:16 | 0.0417 | 0.0418 | 0.0422 | 0.9409 | 0.9409 | 8 | 0.4 |
| **Media concentration 8 g/l** | | | | | | | |
| 16:1 | 1.3352 | 1.3360 | 1.3517 | 0.1079 | 0.1079 | 8 | 0.08 |
| 8:1 | 1.2610 | 1.2618 | 1.2766 | 0.2130 | 0.2130 | 8 | 0.08 |
| 5:1 | 1.1822 | 1.1829 | 1.1969 | 0.3247 | 0.3247 | 8 | 0.08 |
| 3:1 | 1.0640 | 1.0646 | 1.0772 | 0.4922 | 0.4922 | 8 | 0.08 |
| 1:1 | 0.7095 | 0.7100 | 0.7180 | 0.9950 | 0.9950 | 8 | 0.08 |
| 1:3 | 0.3545 | 0.3550 | 0.3590 | 1.4975 | 1.4975 | 8 | 0.08 |
| 1:5 | 0.2364 | 0.2366 | 0.2394 | 1.6649 | 1.6649 | 8 | 0.08 |
| 1:8 | 0.1576 | 0.1577 | 0.1596 | 1.7766 | 1.7766 | 8 | 0.08 |
| 1:16 | 0.0835 | 0.0835 | 0.0845 | 1.8817 | 1.8817 | 8 | 0.08 |
| **Media concentration 20 g/l** | | | | | | | |
| 16:1 | 3.3381 | 3.3401 | 3.3794 | 0.2697 | 0.2697 | 8 | 0.2 |
| 8:1 | 3.1526 | 3.1545 | 3.1916 | 0.5325 | 0.5325 | 8 | 0.2 |
| 5:1 | 2.9556 | 2.9573 | 2.9921 | 0.8117 | 0.8117 | 8 | 0.2 |
| 3:1 | 2.6600 | 2.6615 | 2.6930 | 1.2305 | 1.2305 | 8 | 0.2 |
| 1:1 | 1.7735 | 1.7745 | 1.7955 | 2.4870 | 2.4870 | 8 | 0.2 |
| 1:3 | 0.8865 | 0.8870 | 0.8975 | 3.7435 | 3.7435 | 8 | 0.2 |
| 1:5 | 0.5911 | 0.5915 | 0.5984 | 4.1623 | 4.1623 | 8 | 0.2 |
| 1:8 | 0.3941 | 0.3943 | 0.3990 | 4.4416 | 4.4416 | 8 | 0.2 |
| 1:16 | 0.2086 | 0.2088 | 0.2112 | 4.7044 | 4.7044 | 8 | 0.2 |
| **Media concentration 50 g/l** | | | | | | | |
| 16:1 | 8.3452 | 8.3502 | 8.4484 | 0.6742 | 0.6742 | 8 | 0.5 |
| 8:1 | 7.8816 | 7.8863 | 7.9790 | 1.3312 | 1.3312 | 8 | 0.5 |
| 5:1 | 7.3890 | 7.3934 | 7.4803 | 2.0293 | 2.0293 | 8 | 0.5 |
| 3:1 | 6.6500 | 6.6538 | 6.7325 | 3.0763 | 3.0763 | 8 | 0.5 |
| 1:1 | 4.4331 | 4.4331 | 4.4881 | 6.2175 | 6.2175 | 8 | 0.5 |
| 1:3 | 2.2169 | 2.2181 | 2.2444 | 9.3588 | 9.3588 | 8 | 0.5 |
| 1:5 | 1.4778 | 1.4787 | 1.4961 | 10.406 | 10.406 | 8 | 0.5 |
| 1:6 | 0.9852 | 0.9858 | 0.9974 | 11.104 | 11.104 | 8 | 0.5 |
| 1:16 | 0.5216 | 0.5219 | 0.5280 | 11.761 | 11.761 | 8 | 0.5 |

**Suppl. table S6**

**Supplementary table S6.** Eight-scale categories of fungus colony colour changes in *Metarhizium* fungi as a measure of conidia formation. Example pictures are shown in Supplementary Figure S3.

| Category | Pigmentation |
|---|---|
| 0 | Exploratory growth |
| 1 | White |
| 2 | Light yellow (Beige) |
| 3 | Yellow |
| 4 | Dark yellow (Orange) |
| 5 | Yellow and light green spots |
| 6 | Light green |
| 7 | Green |
| 8 | Dark green (Brownish) |

### Supplementary figures

**Suppl. Figure S1**

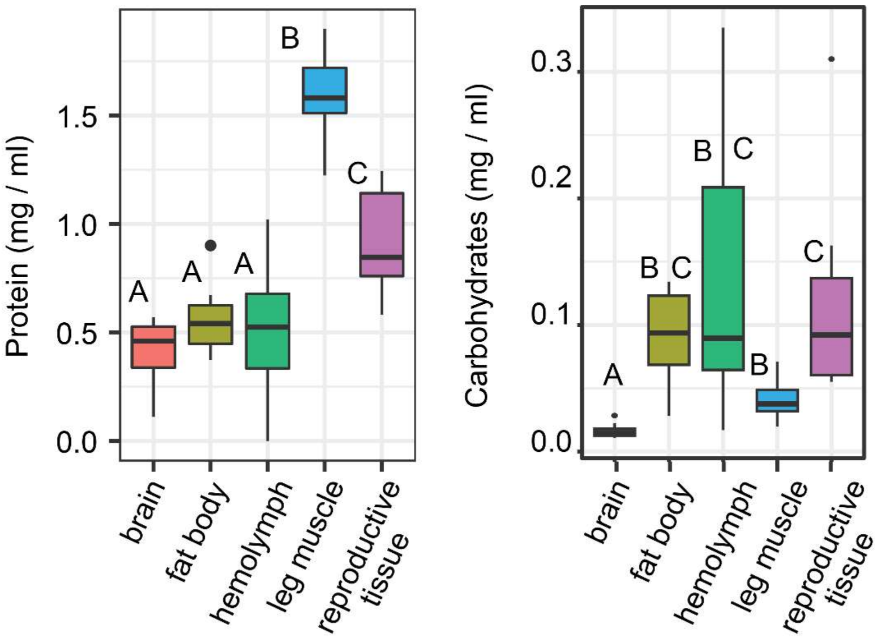


**Suppl. Fig. S1.** Total protein and carbohydrate measurements of specific insect tissues. Letters above each boxplot represents significant group differences following GLM analysis. Solid line inside boxplots represent the median, the box the interquartile 50% of the data, and whiskers represent the distance between maximum and minimum values to the interquartile range, and separate dots are outliers.

**Suppl. Figure S2**

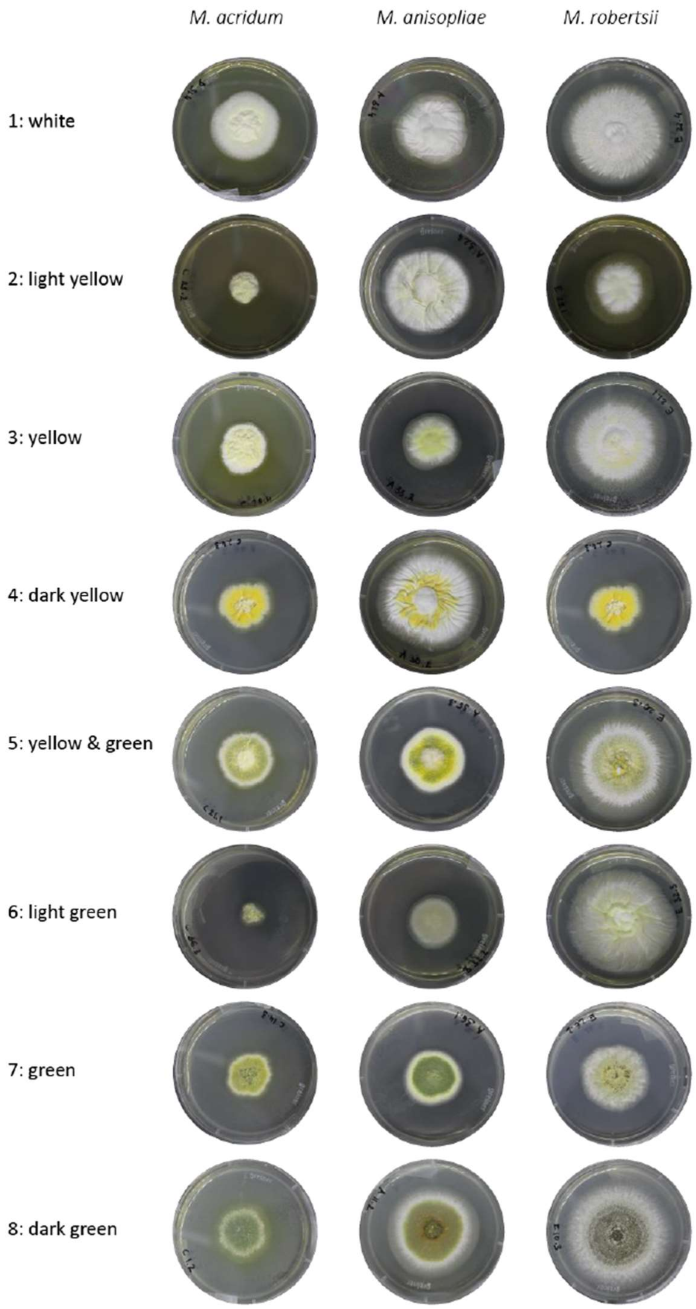


**Suppl. Fig. S2**: Colony pigmentation scoring categories of *Metarhizium* fungi.
