## Supplementary material for "Insect hosts are nutritional landscapes navigated by fungal pathogens": ZIP-file containing data and code: Manuscript_dataAnalysis_211123.html

Data Analysis of De Fine Licht et al.


### Data Analysis of De Fine Licht et al.

#### 2023-11-21

- Load
  packages
- C and P content of Locust
  organs
  - Plot individual organ
    content of C and P
  - Plot RNN of Locust organs
    for C and P
- FNN analysis of fungal
  isolates
  - Statistical
    analysis of individual organ content of C and P
  - FNN statistics per isolate
  - FNN statistics averaged per
    species
- Analysis
  of C and P consumption of fungi in liquid media

### Load packages

```
#Load packages
library(devtools)
library(readxl)
library(tidyverse)
library(ggplot2)
library(nlme)
library(ggpubr)
library(ggrepel)
library(tidyverse)
library(ggConvexHull)
library(car)
library(lme4)
library(plyr)
library(vegan)
library(fields)
library(multcomp)
library(effects)
library(fields)
library(robustHD)
library(utils)
```

### C and P content of Locust organs

#### Plot individual organ content of C and P

```
one <- read_excel("organ_pc_r.xlsx", sheet = "Sheet1")

attach(one)
# grouped boxplot

C2 <- ggplot(one, aes(x=tissue, y=mean_prot_mg_ml, fill=tissue)) + 
  geom_boxplot()+
  theme_bw(base_size = 16) +
  theme(axis.text.x = element_text(angle = 90, hjust = 1))

C2

V2 <- ggplot(one, aes(x=tissue, y=mean_carb_mg_ml, fill=tissue)) + 
  geom_boxplot()+
  theme_bw(base_size = 16) +
  theme(axis.text.x = element_text(angle = 90, hjust = 1))

V2

# plot side by side
require(gridExtra)
grid.arrange(C2, V2, ncol=2)
```

#### Plot RNN of Locust organs for C and P

```
ggplot(DF, aes(x = tissue, y = Value))+
  geom_boxplot()+
  facet_wrap(.~Variable1, scales = "free")+
  theme(axis.text.x = element_text(angle = 45, hjust = 1))+


r1 +  theme_classic() + scale_x_continuous(expand = c(0, 0), limits = c(0, 2)) + 
  scale_y_continuous(expand = c(0, 0), limits = c(0, 0.5)) + coord_fixed() +
  geom_polygon(data = hull, alpha = 0.2,
               aes(fill = tissue,colour = tissue))+
  labs(x = "protein mg/mL", y = "carbohydrate mg/mL") + theme(aspect.ratio = 1)


#final RNN plots

r1 <- ggplot(one, aes(x = mean_prot_mg_ml, y = mean_carb_mg_ml, color = factor(tissue))) +
  geom_point(aes(color=factor(tissue))) 

hull <- one %>% group_by(tissue) %>%
  slice(chull(mean_prot_mg_ml, mean_carb_mg_ml))

myplot <- r1 +  theme_classic() + scale_x_continuous(expand = c(0, 0), limits = c(0, 2)) + 
  scale_y_continuous(expand = c(0, 0), limits = c(0, 0.4)) + coord_fixed() +
  geom_polygon(data = hull, alpha = 0.2,
               aes(fill = tissue,colour = tissue))+
  labs(x = "Protein mg/ml", y = "Carbohydrates mg/ml") + theme(aspect.ratio = 1)

print(myplot)
```

### FNN analysis of fungal isolates

#### Statistical analysis of individual organ content of C and P

```
#what are organ pct p and pct c concentrations
attach(one)

summary(mean_prot_mg_ml)
summary(mean_carb_mg_ml)


#STEP 1: normality test
#(Visual Method) Create a histogram.

attach(a)
hist(mean_prot_mg_ml, col='steelblue', main='Normal test')
hist(mean_carb_mg_ml, col='steelblue', main='Normal test')


#(Visual Method) Create a Q-Q plot.

qqnorm(mean_prot_mg_ml, main='Normal')
qqline(mean_prot_mg_ml)

qqnorm(mean_carb_mg_ml, main='Normal')
qqline(mean_carb_mg_ml)


#(Formal Statistical Test) Perform a Shapiro-Wilk Test.

shapiro.test(mean_prot_mg_ml)
shapiro.test(mean_carb_mg_ml)

#because of significant shapiro wilks test, using non-parametric 
#test to test signifcant Do organs have different protein and/or carb amounts?

kruskal.test( mean_prot_mg_ml ~ tissue, data = one)
pairwise.wilcox.test(mean_prot_mg_ml, tissue,
                     p.adjust.method = "bonferroni")
pairwise.wilcox.test(mean_prot_mg_ml, tissue,
                     p.adjust.method = "BH")

kruskal.test( mean_carb_mg_ml ~ tissue, data = one)
pairwise.wilcox.test(mean_carb_mg_ml, tissue,
                     p.adjust.method = "bonferroni")
pairwise.wilcox.test(mean_carb_mg_ml, tissue,
                     p.adjust.method = "BH")

#did organs have more protein than carbohydrates?

t.test(mean_prot_mg_ml, mean_carb_mg_ml, paired = TRUE, alternative = "two.sided")
wilcox.test(mean_prot_mg_ml, mean_carb_mg_ml, paired = TRUE, alternative = "two.sided")
wilcox.test(mean_prot_mg_ml, mean_carb_mg_ml, paired = TRUE, exact=TRUE)
```

#### FNN statistics per isolate

```
#FNN analyses

a <- read_excel("master_fnn.xlsx", sheet = "avg_fnn")
a

b <- read_excel("master_fnn.xlsx", sheet = "isolate_fnn")
b

#FNN stats: individual isolate analyses (dataset b)
attach(b)
# each isolate individually
#A_anis: M. anisopliae KVL  18_19 (ESALQ_1116)
#B_anis: M. anisopliae KVL  18_26 (ESALQ_1604)
#A1_anis: M. anisopliae KVL  18_05 (ARSEF_549)
#C_acri: M. acridum KVL 04_55 (ARSEF_7486)
#D_acri: M. acridum KVL  18_02 (ARSEF_3609)
#B1_acri: M. acridum KVL  12_13 (ARSEF_324)
#E_rob: M. robertsii KVL  12_35 (KVL_12-35)
#F_rob M. robertsii KVL  13_12 (KVL_13-12)


A_anis<-subset(b, isolate=="A_anis")
A1_anis<-subset(b, isolate=="A1_anis")
B_anis<-subset(b, isolate=="B_anis")
B1_acri<-subset(b, isolate=="B1_acri")
C_acri<-subset(b, isolate=="C_acri")
D_acri<-subset(b, isolate=="D_acri")
E_rob<-subset(b, isolate=="E_rob")
F_rob<-subset(b, isolate=="F_rob")


# area
model.A_anis<-lm(area ~ pct_prot + pct_carb + pct_prot:pct_carb + I(pct_prot^2) + I(pct_carb^2), data=A_anis)
summary(model.A_anis)
hist(resid(model.A_anis))
Anova(model.A_anis)

model.A1_anis<-lm(area ~ pct_prot + pct_carb + pct_prot:pct_carb + I(pct_prot^2) + I(pct_carb^2), data=A1_anis)
summary(model.A1_anis)
hist(resid(model.A1_anis))
Anova(model.A1_anis)

model.B_anis<-lm(area ~ pct_prot + pct_carb + pct_prot:pct_carb + I(pct_prot^2) + I(pct_carb^2), data=B_anis)
summary(model.B_anis)
hist(resid(model.B_anis))
Anova(model.B_anis)

model.B1_acri<-lm(area ~ pct_prot + pct_carb + pct_prot:pct_carb + I(pct_prot^2) + I(pct_carb^2), data=B1_acri)
summary(model.B1_acri)
hist(resid(model.B1_acri))
Anova(model.B1_acri)

model.C_acri<-lm(area ~ pct_prot + pct_carb + pct_prot:pct_carb + I(pct_prot^2) + I(pct_carb^2), data=C_acri)
summary(model.C_acri)
hist(resid(model.C_acri))
Anova(model.C_acri)

model.D_acri<-lm(area ~ pct_prot + pct_carb + pct_prot:pct_carb + I(pct_prot^2) + I(pct_carb^2), data=D_acri)
summary(model.D_acri)
hist(resid(model.D_acri))
Anova(model.D_acri)

model.E_rob<-lm(area ~ pct_prot + pct_carb + pct_prot:pct_carb + I(pct_prot^2) + I(pct_carb^2), data=E_rob)
summary(model.E_rob)
hist(resid(model.E_rob))
Anova(model.E_rob)

model.F_rob<-lm(area ~ pct_prot + pct_carb + pct_prot:pct_carb + I(pct_prot^2) + I(pct_carb^2), data=F_rob)
summary(model.F_rob)
hist(resid(model.F_rob))
Anova(model.F_rob)

#color
model.A_anis<-lm(color ~ pct_prot + pct_carb + pct_prot:pct_carb + I(pct_prot^2) + I(pct_carb^2), data=A_anis)
summary(model.A_anis)
hist(resid(model.A_anis))
Anova(model.A_anis)

model.A1_anis<-lm(color ~ pct_prot + pct_carb + pct_prot:pct_carb + I(pct_prot^2) + I(pct_carb^2), data=A1_anis)
summary(model.A1_anis)
hist(resid(model.A1_anis))
Anova(model.A1_anis)

model.B_anis<-lm(color ~ pct_prot + pct_carb + pct_prot:pct_carb + I(pct_prot^2) + I(pct_carb^2), data=B_anis)
summary(model.B_anis)
hist(resid(model.B_anis))
Anova(model.B_anis)

model.B1_acri<-lm(color ~ pct_prot + pct_carb + pct_prot:pct_carb + I(pct_prot^2) + I(pct_carb^2), data=B1_acri)
summary(model.B1_acri)
hist(resid(model.B1_acri))
Anova(model.B1_acri)

model.C_acri<-lm(color ~ pct_prot + pct_carb + pct_prot:pct_carb + I(pct_prot^2) + I(pct_carb^2), data=C_acri)
summary(model.C_acri)
hist(resid(model.C_acri))
Anova(model.C_acri)

model.D_acri<-lm(color ~ pct_prot + pct_carb + pct_prot:pct_carb + I(pct_prot^2) + I(pct_carb^2), data=D_acri)
summary(model.D_acri)
hist(resid(model.D_acri))
Anova(model.D_acri)

model.E_rob<-lm(color ~ pct_prot + pct_carb + pct_prot:pct_carb + I(pct_prot^2) + I(pct_carb^2), data=E_rob)
summary(model.E_rob)
hist(resid(model.E_rob))
Anova(model.E_rob)

model.F_rob<-lm(color ~ pct_prot + pct_carb + pct_prot:pct_carb + I(pct_prot^2) + I(pct_carb^2), data=F_rob)
summary(model.F_rob)
hist(resid(model.F_rob))
Anova(model.F_rob)

#spore (log10spore)
model.A_anis<-lm(spore ~ pct_prot + pct_carb + pct_prot:pct_carb + I(pct_prot^2) + I(pct_carb^2), data=A_anis)
summary(model.A_anis)
hist(resid(model.A_anis))
Anova(model.A_anis)

model.A1_anis<-lm(spore ~ pct_prot + pct_carb + pct_prot:pct_carb + I(pct_prot^2) + I(pct_carb^2), data=A1_anis)
summary(model.A1_anis)
hist(resid(model.A1_anis))
Anova(model.A1_anis)

model.B_anis<-lm(spore ~ pct_prot + pct_carb + pct_prot:pct_carb + I(pct_prot^2) + I(pct_carb^2), data=B_anis)
summary(model.B_anis)
hist(resid(model.B_anis))
Anova(model.B_anis)

model.B1_acri<-lm(spore ~ pct_prot + pct_carb + pct_prot:pct_carb + I(pct_prot^2) + I(pct_carb^2), data=B1_acri)
summary(model.B1_acri)
hist(resid(model.B1_acri))
Anova(model.B1_acri)

model.C_acri<-lm(spore ~ pct_prot + pct_carb + pct_prot:pct_carb + I(pct_prot^2) + I(pct_carb^2), data=C_acri)
summary(model.C_acri)
hist(resid(model.C_acri))
Anova(model.C_acri)

model.D_acri<-lm(spore ~ pct_prot + pct_carb + pct_prot:pct_carb + I(pct_prot^2) + I(pct_carb^2), data=D_acri)
summary(model.D_acri)
hist(resid(model.D_acri))
Anova(model.D_acri)

model.E_rob<-lm(spore ~ pct_prot + pct_carb + pct_prot:pct_carb + I(pct_prot^2) + I(pct_carb^2), data=E_rob)
summary(model.E_rob)
hist(resid(model.E_rob))
Anova(model.E_rob)

model.F_rob<-lm(spore ~ pct_prot + pct_carb + pct_prot:pct_carb + I(pct_prot^2) + I(pct_carb^2), data=F_rob)
summary(model.F_rob)
hist(resid(model.F_rob))
Anova(model.F_rob)
```

#### FNN statistics averaged per species

```
#FNN stats: mean species analyses (dataset a)
attach(a)
# each isolate individually
av_acri<-subset(a, sp=="av_acri")
av_anis<-subset(a, sp=="av_anis")
av_ef_rob<-subset(a, sp=="av_ef_rob")

# area AVG sp
model.av_acri<-lm(area ~ pct_prot + pct_carb + pct_prot:pct_carb + I(pct_prot^2) + I(pct_carb^2), data=av_acri)
summary(model.av_acri)
hist(resid(model.av_acri))
Anova(model.av_acri)

model.av_anis<-lm(area ~ pct_prot + pct_carb + pct_prot:pct_carb + I(pct_prot^2) + I(pct_carb^2), data=av_anis)
summary(model.av_anis)
hist(resid(model.av_anis))
Anova(model.av_anis)

model.av_ef_rob<-lm(area ~ pct_prot + pct_carb + pct_prot:pct_carb + I(pct_prot^2) + I(pct_carb^2), data=av_ef_rob)
summary(model.av_ef_rob)
hist(resid(model.av_ef_rob))
Anova(model.av_ef_rob)

# color AVG sp
model.av_acri<-lm(color ~ pct_prot + pct_carb + pct_prot:pct_carb + I(pct_prot^2) + I(pct_carb^2), data=av_acri)
summary(model.av_acri)
hist(resid(model.av_acri))
Anova(model.av_acri)

model.av_anis<-lm(color ~ pct_prot + pct_carb + pct_prot:pct_carb + I(pct_prot^2) + I(pct_carb^2), data=av_anis)
summary(model.av_anis)
hist(resid(model.av_anis))
Anova(model.av_anis)

model.av_ef_rob<-lm(color ~ pct_prot + pct_carb + pct_prot:pct_carb + I(pct_prot^2) + I(pct_carb^2), data=av_ef_rob)
summary(model.av_ef_rob)
hist(resid(model.av_ef_rob))
Anova(model.av_ef_rob)


# spore AVG sp
model.av_acri<-lm(spore ~ pct_prot + pct_carb + pct_prot:pct_carb + I(pct_prot^2) + I(pct_carb^2), data=av_acri)
summary(model.av_acri)
hist(resid(model.av_acri))
Anova(model.av_acri)

model.av_anis<-lm(spore ~ pct_prot + pct_carb + pct_prot:pct_carb + I(pct_prot^2) + I(pct_carb^2), data=av_anis)
summary(model.av_anis)
hist(resid(model.av_anis))
Anova(model.av_anis)

model.av_ef_rob<-lm(spore ~ pct_prot + pct_carb + pct_prot:pct_carb + I(pct_prot^2) + I(pct_carb^2), data=av_ef_rob)
summary(model.av_ef_rob)
hist(resid(model.av_ef_rob))
Anova(model.av_ef_rob)


#HEAT MAPS fields export as pdf - example for one map provided
#isolate mean area across percent P and C
require(fields)
attach(b)

pdf(file="A_anis_area.pdf")
fit = Tps(cbind(pct_prot,pct_carb), area, lambda=0.0005, data=A_anis)
surface(fit, type="C",xlab="Protein", ylab="Carbohydrate", nx=200,ny=200, lab=c(6,6,6), font.lab=1, font.axis=1, labcex=1.4, cex.axis=1.4, cex.lab=1.4, xlim=c(0,80), ylim=c(0,80))
dev.off()


#isolate mean staph dens across percent P and C
require(fields)
attach(e)
pdf(file="meanAE_staphdens_pctPC.pdf")
fit = Tps(cbind(pct_prot,pct_carb), staphdens, lambda=0.001)
surface(fit, type="C",xlab="Protein", ylab="Carbohydrate", nx=200,ny=200, lab=c(6,6,6), font.lab=1, font.axis=1, labcex=1.4, cex.axis=1.4, cex.lab=1.4, xlim=c(0,80), ylim=c(0,80))
dev.off()
```

### Analysis of C and P consumption of fungi in liquid media

```
# Read the data
protcarb<-read.csv2("ProtCarb_consump_data.csv")

# check the file
head(protcarb)
str(protcarb)

# Subset the data into files for each fungal species
# A = M. anisopliae
# B = M. acridum
# C = M. robertsii
Afungus <- subset(protcarb, Fungus=="A")
Bfungus <- subset(protcarb, Fungus=="B")
Cfungus <- subset(protcarb, Fungus=="C")

# subset per media composition
Afungus_d50_13 <- subset(Afungus,media=="d50_13")
Afungus_d50_31 <- subset(Afungus,media=="d50_31")
Afungus_d15_13 <- subset(Afungus,media=="d15_13")
Afungus_d15_31 <- subset(Afungus,media=="d15_31")

# Define the models
ModelA_d50_13 <- lm(carb ~ prot, data=Afungus_d50_13)
ModelA_d50_31 <- lm(carb ~ prot, data=Afungus_d50_31)
ModelA_d15_13 <- lm(carb ~ prot, data=Afungus_d15_13)
ModelA_d15_31 <- lm(carb ~ prot, data=Afungus_d15_31)

# Have a look at the estimated coefficients of the linear model
coef(ModelA_d50_31)
coef(ModelA_d50_13)
coef(ModelA_d15_31)
coef(ModelA_d15_13)

# test the models comparing them to the ratio of protein and carbohydrate in
# the media

# ModelA_d50_31
linearHypothesis(ModelA_d50_31, "prot = 0.33")
# ModelA_d50_13
linearHypothesis(ModelA_d50_13, "prot = 3.00")
# ModelA_d15_31
linearHypothesis(ModelA_d15_31, "prot = 0.33")
# ModelA_d50_13
linearHypothesis(ModelA_d15_13, "prot = 3.00")

Bfungus_d50_13 <- subset(Bfungus,media=="d50_13")
Bfungus_d50_31 <- subset(Bfungus,media=="d50_31")
Bfungus_d15_13 <- subset(Bfungus,media=="d15_13")
Bfungus_d15_31 <- subset(Bfungus,media=="d15_31")

ModelB_d50_13 <- lm(carb ~ prot, data=Bfungus_d50_13)
ModelB_d50_31 <- lm(carb ~ prot, data=Bfungus_d50_31)
ModelB_d15_13 <- lm(carb ~ prot, data=Bfungus_d15_13)
ModelB_d15_31 <- lm(carb ~ prot, data=Bfungus_d15_31)

coef(ModelB_d50_31)
coef(ModelB_d50_13)
coef(ModelB_d15_31)
coef(ModelB_d15_13)
# ModelB_d50_31
linearHypothesis(ModelB_d50_31, "prot = 0.33")
# ModelB_d50_13
linearHypothesis(ModelB_d50_13, "prot = 3.00")
# ModelB_d15_31
linearHypothesis(ModelB_d15_31, "prot = 0.33")
# ModelB_d15_13
linearHypothesis(ModelB_d15_13, "prot = 3.00")

Cfungus_d50_13 <- subset(Cfungus,media=="d50_13")
Cfungus_d50_31 <- subset(Cfungus,media=="d50_31")
Cfungus_d15_13 <- subset(Cfungus,media=="d15_13")
Cfungus_d15_31 <- subset(Cfungus,media=="d15_31")

ModelC_d50_13 <- lm(carb ~ prot, data=Cfungus_d50_13)
ModelC_d50_31 <- lm(carb ~ prot, data=Cfungus_d50_31)
ModelC_d15_13 <- lm(carb ~ prot, data=Cfungus_d15_13)
ModelC_d15_31 <- lm(carb ~ prot, data=Cfungus_d15_31)

coef(ModelC_d50_31)
coef(ModelC_d50_13)
coef(ModelC_d15_31)
coef(ModelC_d15_13)
# ModelC_d50_31
linearHypothesis(ModelC_d50_31, "prot = 0.33")
# ModelC_d50_13
linearHypothesis(ModelC_d50_13, "prot = 3.00")
# ModelC_d15_31
linearHypothesis(ModelC_d15_31, "prot = 0.33")
# ModelC_d15_13
linearHypothesis(ModelC_d15_13, "prot = 3.00")
```
